# Genome-wide, Organ-delimited gene regulatory networks (OD-GRNs) provide high accuracy in candidate TF selection across diverse processes

**DOI:** 10.1101/2023.06.17.542927

**Authors:** Rajeev Ranjan, Sonali Srijan, Somaiah Balekuttira, Tina Agarwal, Melissa Ramey, Madison Dobbins, Xiaojin Wang, Karen Hudson, Ying Li, Kranthi Varala

## Abstract

Construction of organ-specific gene expression datasets that include hundreds to thousands of experiments would greatly aid reconstruction of gene regulatory networks with organ-level spatial resolution. However, creating such datasets is greatly hampered by the requirements of extensive and tedious manual curation. Here we trained a supervised classification model that can accurately classify the organ-of-origin for a plant transcriptome. This K-Nearest Neighbor-based multiclass classifier was used to create organ-specific gene expression datasets for the leaf, root, shoot, flower, seed, seedling, silique, and stem in the model plant *Arabidopsis thaliana*. In the leaf, root, flower, seed and, a gene regulatory network (GRN) inference approach was used to determine: *i*. influential transcription factors (TFs) in that organ and, *ii*. the most influential TFs for specific biological processes in the organ. These genome-wide, organ-delimited GRNs (OD-GRNs), identified *de novo* many known regulators of organ development and processes operating in those organs. Moreover, many previously unknown TF regulators were highly ranked as potential master regulators of organ development or organ-specific processes. As a proof-of-concept, we focused on experimentally validating the predicted TF regulators of lipid biosynthesis in seeds, with relevance to food and biofuel production. Of the top twenty candidate TFs, eight (e.g., WRI1, LEC1, and FUS3) are known regulators of seed oil content. Importantly, we validated that seven more candidate TFs, whose role was previously unknown in seed lipid biosynthesis, indeed affect this process by genetics and physiological approaches, thus yielding a net accuracy rate of >75% for the *de novo* TF predictions. The general approach developed here could be extended to any species with sufficiently large gene expression datasets to speed up hypothesis generation and testing for constructing gene regulatory networks at a high spatial resolution.

**Significance Statement:** Our study develops a machine-learning framework for building extremely large gene expression datasets for each organ, and to infer organ-delimited gene regulatory networks. We show that this approach is very successful at predicting which transcription factors are going to regulate processes at an organ level. We validated the accuracy of the predictions for transcription factor regulators using the seed lipid synthesis pathway as a case study. We demonstrated a very high success rate for uncovering both known and novel transcription factor regulators for the seed lipid biosynthesis pathway. The approach described in this study is broadly applicable across any organism (plant or animal) that has a large body of public gene expression data.

## INTRODUCTION

An organism’s gene expression profile is modulated in response to its environment, primarily through the action of a special class of genes called transcription factors (TFs). Therefore, an important goal of molecular biology has been the discovery of transcription factor (TF) to target gene interactions, which at a genome-wide scale are called gene regulatory networks (GRNs). However, the inference of GRNs remains a challenge (1) due to the high variability in gene expression and the presence of transient, context-dependent TF-target relationships, which results in low signal-to-noise ratio. With the advent of cheap and robust mRNA sequencing technologies, thousands of independent mRNA assays in model organisms has resulted in tens of thousands of unique genome-wide expression assays. This massive dataset may be used to infer the GRNs provided two challenges can be surmounted: 1. Computational cost of working with these massive data sets and 2. Coherence on interactions across development, growth and treatment variations. In plants, numerous studies (2) have focused on the inference of GRNs from gene expression data. The model organism *Arabidopsis thaliana* is well suited for genome-wide GRN reconstruction for the following reasons: *1*. compact genome encodes approximately 28,000 genes which includes ∼ 1,700 TFs (i.e., ∼30 million potential regulatory interactions), *2*. massive gene expression dataset with tens of thousands of individual assays and *3*. modest organismal complexity with few distinct organs (e.g., a shoot made primarily of leaves, simple flowers etc.)

Individual TFs can potentially bind (3) and/or regulate (4) hundreds to thousands of targets. However, the precise targets of a TF in a given organ and/or condition seems to be constrained by promoter availability (5) and likely other factors. The most obvious delineation of GRN composition and this global gene expression is the plant organ (6–8). Traditionally, GRN inference has worked best on large expression sets from the same organ after an application of one or more external signals (5, 9, 10). While this approach is powerful in discovering key TF regulators of plant responses, its application across multiple responses is limited by the cost and labor-intensive sampling required to conduct all the required experiments, perform gene expression assays, and the computational cost to process them. Thus, it is beneficial to precompute genome-wide GRNs from public gene expression datasets that include repeated measurements of most, if not all, the genes functional in an organ.

The focus of our study is to enable such GRN construction by: *i*. Building a comprehensive gene expression dataset, *ii*. Use an automated classifier to create unbiased organ delineated data sets, *iii*. Infer GRNs functional in each organ and *iv*. Assay the accuracy of this approach by functional validation of predicted regulator roles. As a proof-of-concept, we chose to focus on the process of lipid biosynthesis in the seed. Substantial efforts in understanding the synthesis, storage and degradation of oils have led to a well-characterized biochemical pathway (11). Significant attention has also been paid to the transcriptional regulation of these lipid biosynthesis genes leading to the discovery of a handful of TFs that regulate this process (e.g., WRI1, LEC1, FUS3 etc.)(12). WRINKLED1 (WRI1) is a master regulator of seed oil accumulation as it regulates many genes involved in glycolysis and fatty acid (FA) biosynthesis (13–15). LEC1, LEC2, FUS3, and ABI3, are other TFs whose roles have been established in seed oil biosynthesis (16–20). Among them, LEC1, LEC2, FUS3, and ABI3 work upstream of WRI1 and together with HSI2/VAL1, AGL15, BBM, TT8, and MYB89 form a network that tightly controls the expression of WRI1(15–17, 21, 22). Additionally, the TFs MYB96, TZF, WRKY10, WRKY43, and bZIP67 positively regulate oil accumulation in seeds while, TT2, TTG1, GL2, MYB76, MYB118, and WRKY6, negatively affect seed oil content (23– 32). Despite this wealth of knowledge, robust strategies to increase the overall content and especially the composition of storage lipids are somewhat lacking. Given the size and complexity of this pathway (over 100 genes, spread across multiple organelles) and the central role it plays in seed development, it is likely that the transcriptional regulation of the pathway is complex and includes additional, as yet uncharacterized, TFs. Therefore, we chose the discovery and validation of TFs regulating the lipid metabolism pathway in the seeds as a target to assess the accuracy of our approach in identifying TFs that regulate a pathway or process of interest. A successful inference strategy would identify *de novo* the known regulators as well as predict unknown TF regulators of seed lipid biosynthesis.

## RESULTS

### Compiling a large, labeled gene expression dataset to enable machine learning

*Arabidopsis thaliana* as a model organism has been subjected to gene expression assays for decades using a variety of platforms. For this study, we chose to select the largest subset of experiments that used the Ilumina platform to assay expression via RNA-Seq from the NCBI’s short read archive (SRA) database. This public repository was queried in Fall 2018, which generated a data set of 18,174 individual Illumina runs (Supp. Table 1). A quasi mapping algorithm (33) was used to convert raw sequence data into a gene expression estimate for 16,830 runs after eliminating experiments with too few mappable reads (see Methods). The gene expression matrix was subjected to a full quantile normalization (34) and transformed to a log2 scale.

Concurrently, we collected the metadata associated with each experiment from the SRA database. The organ from which the RNA was extracted from, *i*.*e*. the label, was coded by manual curation of the metadata. In cases where organ assignment was not provided in the metadata, but the experiment was associated with a published manuscript, the organ-of-origin was retrieved from the corresponding publication. In this way, a total of nine organ labels, including Aerial, Flower, Leaf, Root, Seed, Seedling, Shoot, Silique, and Stem, were created and assigned to the 16,830 RNAseq runs. Therefore, we compiled a fully labeled, high confidence gene expression dataset that is ideally suited to build a machine learning (ML) model in a supervised learning approach. Technically, each data point (i.e. individual RNAseq data) in this set includes 33,584 featuresin the Arabidopsis genome (35), as well as and one of the nine organ labels.

To capture each organ’s global expression in all the stages/conditions they have been sampled in various experiments, we didn’t apply any filters to select a specific condition, developmental stage, or environmental treatment. Given this diversity of conditions, it is not apparent that all experiments from the same organ would cluster together and away from the other organ clusters. To verify whether the organ-of-origin is the major determinant of global gene expression, we plotted all experiments into a single two-dimensional space, using the tSNE dimensionality reduction approach. As shown in Figure 1A, most experiments from the same organ cluster together to form a distinct global profile for each organ (as indicated by color in Fig. 1A). However, it is also apparent that the same organ can exist in multiple stable states that differ substantially from each other, as is seen in the case of Leaf samples (multiple clusters of green dots) and the three distinct clusters formed by the root samples (brown dots). This suggests that a simple clustering approach (e.g., K-Means) is unlikely to create clean organ-specific subsets from this large set. However, machine learning approaches have proven extremely robust in such supervised classification problems in biology (36, 37).

**Figure 1.**
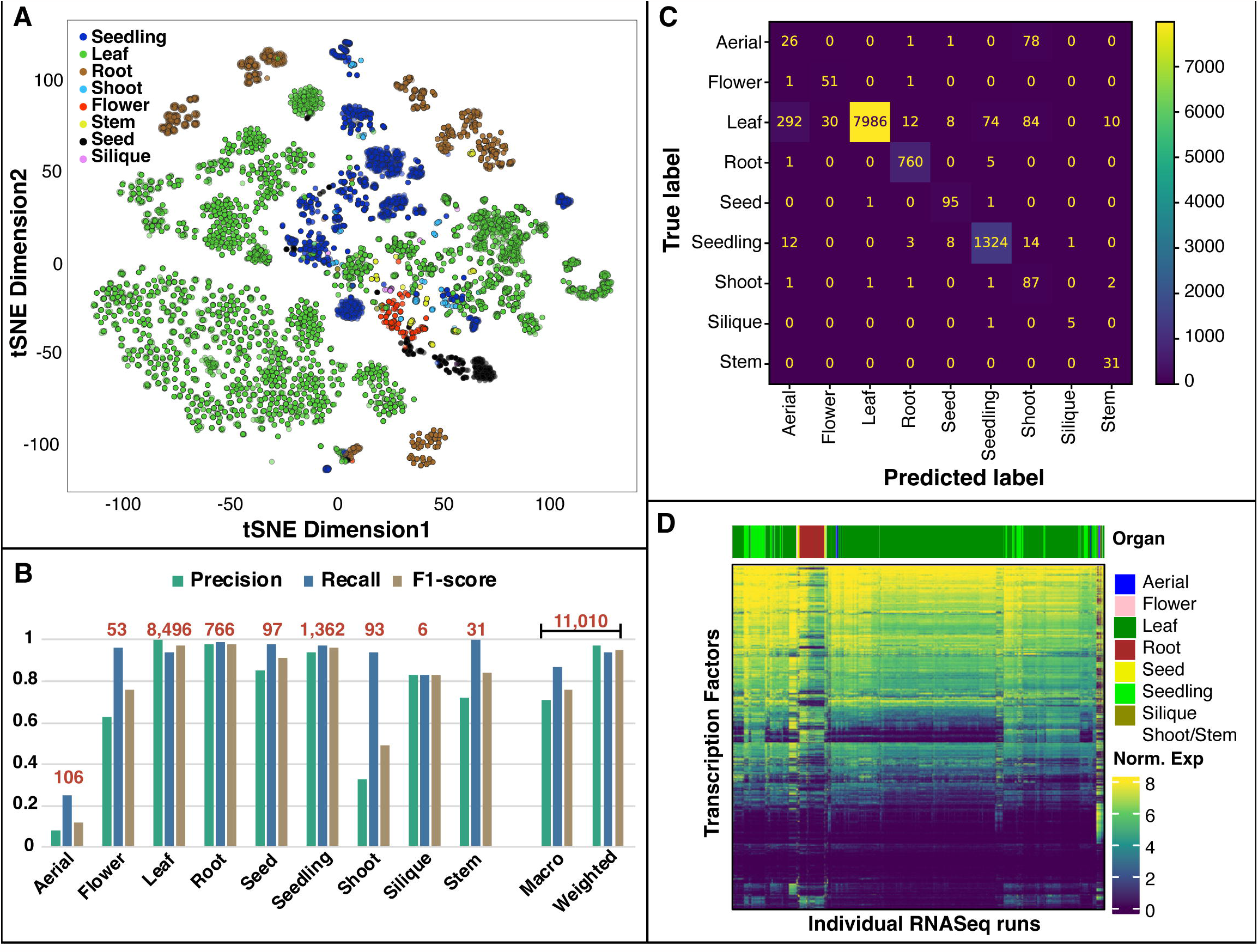
ML classifier creates organ-specific gene expression datasets. **A**. Organs are the main determinants of global gene expression as demonstrated by the strong organ driven clustering in this reduced dimensionality (tSNE) plot. **B**. KNN-based classifier achieved high accuracy (weighted F1 =0.95). Precision, recall, F1) are co-variate and related to the number of samples in the training set (Fig. 1 B, numbers in red). **C**. Confusion matrix of true vs. predicted labels from the KNN-based classifier shows that the majority of mislabeling is due to biological ambiguity of organ labels. **D**. Most TFs are expressed across many organs (normalized expression shown).

### Supervised classifier assigns organ labels with high accuracy

We explored three machine learning models to determine the best performing classifier, i.e., one with high precision and recall. A K-Nearest-Neighbor (KNN) based, a Support Vector Machine (SVM) based, and a Neural Network (NN) based model was each trained using 90:10% split of training and testing data and a cross-validation strategy (see Methods for details). The NN model performed poorest with both precision and accuracy scores well below the KNN and SVM models across all organs, except the leaf (data not shown). Between the KNN and SVM based models, the KNN model had higher precision and recall and F1 scores across multiple organs (Supp. Fig 1). Therefore, we selected the KNN based classifier as our best performing model. This classifier can determine the correct organ label for most data (macro F1=0.76, weighted F1=0.95; macro F1 is the unweighted average of F1 values across all classes, weighted F1 assigns higher weight to larger classes (no.of true labels) i.e., it adjusts for the size of the class) (Fig. 1B, Supp. Table 2). The performance of the classifier for each organ is directly linked to the amount of available data sets in the training sample as well as the biological ambiguity of organ definition (Fig. 1C). For example, the classifier has nearly perfect precision and accuracy for the leaf which is a clearly demarcated organ and has the largest training set (Fig. 1B&C). The confusion matrix (Fig. 1C) shows that the misclassification of datasets is often due to organ ambiguity. For example, the most common label for false negatives (FN) for the leaf is Aerial, which is an ambiguous “organ” definition in Arabidopsis that is primarily made of leaves. Conversely, the most common true label for the false positives (FP) for the shoot is the leaf which is part of the shoot. Outside of this biological ambiguity, the classifier demonstrated a high degree of accuracy.

It would be very beneficial to GRN inference and a host of other gene expression-based applications if high precision organ sets can be constructed in an unbiased and robust manner in plant and animal species. To assay whether such classifiers can be readily built in other species, we investigated the feasibility of building similar classifiers in tomato. A set of approximately 3,000 RNA-Seq studies in tomato were expertly curated to assign the correct organ label. An xgboost based classifier was trained in tomato and again achieved high accuracy for all organs with at least 100 samples in the training set (precision and recall > 0.9, Supp. Fig 3).

Since, TFs are the primary gene expression regulators, we investigated whether TFs themselves are expressed in an organ specific manner using the Tau metric (38). Briefly, the Tau metric assigns a numeric score between 0-1 to each gene based on how biased its expression is skewed to one or more organs (see Supp. Fig 2). The expression patterns in Fig. 1D and the low Tau scores (Supp. Fig 2), demonstrate that most TFs are not expressed in an organ specific manner. Thus, it is likely that the function of a TF, i.e., the set of genes it regulates, needs to be studied narrowly within the context of the organ, and not broadly at an organism level.

### Inferring organ-delimited gene regulatory networks (OD-GRNs) predicts regulators for an organ, a specific biological process, or a gene-set-of-interest

We hypothesized that large expression sets from a single organ would encode non-conflicting TF-target regulation signals and that using these high-quality sets for GRN inference would result in high accuracy in predicting the roles of TFs. To test these hypotheses, we inferred GRNs in five of the nine organs identified independently to determine the set of TFs that act as primary regulators of a given process within the context of that organ. The five organs chosen due to availability of sufficient data (n>100) as well as their distinct biological definition were: leaf, root, shoot, seed and flower. Further, a random set of 1,000 experiments were chosen from the leaf to limit computational costs. A random forest based network inference tool for TF-target interactions (39), called GENIE3 (40), was used for GRN inference using all 1,717 TFs (41) encoded in the Arabidopsis genome as predictors. In each organ, the top 10% scoring TF-target predictions were retained to create a high confidence organ-delimited GRN (OD-GRN). that includes regulatory predictions for most genes and TFs in an organ. These OD-GRNs may be visualized as a network where both the TFs and genes are nodes connected by directed edges from a TF to its target genes. Each edge has a score that depicts the confidence of the prediction. The net effect of each TF in the OD-GRN was then measured as the sum of scores for all outgoing edges from that TF to generate a ranked list of TFs (Fig. 2A). The top 100 TFs in each OD-GRN were compared to assay the relative importance of TFs across organs. The top TFs in each organ are mostly unique with only a handful of TFs ranking high across organs (Supp. Figure 4). Further, the expression pattern of the union set of top 10 TFs in each organ shows that the TFs with a impactful role in each organ are largely expressed across multiple organs (Supp. Figure 5).

**Figure 2.**
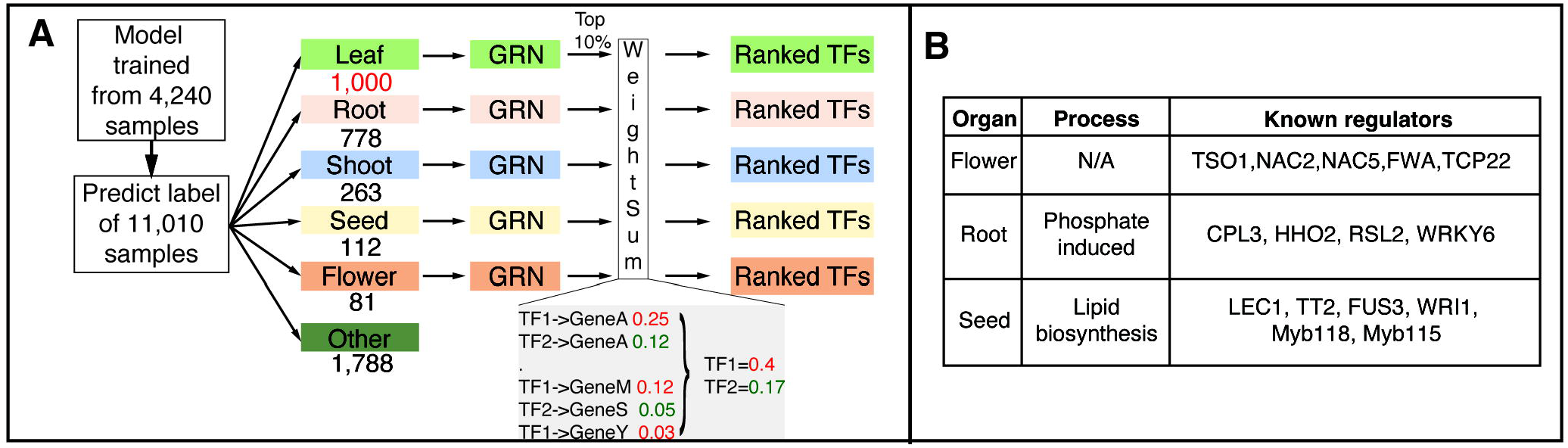
Organ-delimited GRN inference and TF ranking recovers known and predicts candidate regulators of diverse biological processes. **A**. Leaf, root, shoot, seed and flower expression sets were used to infer their GRNs. In each organ the influence of every TF is summarized as the sum of weights of all its edges **B**. Validity of this approach is established by the de novo re-discovery of multiple known regulators of floral development, phosphate induced genes, and lipid biosynthesis genes in the flower, root and seed GRNs respectively.

The OD-GRNs constructed here can be queried repeatedly using subsets of genes (e.g. genes associated with a particular biological process) to generate subnetworks. Each subnetwork will have its own ranked TF lists where the highest ranked TFs are presumed to be most influential on the target genes used to query the OD-GRN. To evaluate the effectiveness of this GRN inference approach to the discovery of TF regulators, we investigated three use cases at the *i*. organ, *ii. gene set* and *iii*. process subnetwork levels. We hypothesize that given a cohesive set of genes serve as bait for network query, the most highly ranked TFs predicted to regulate them should *de novo* discover validated regulators of that gene set as well as predict novel ones.

#### Case study 1: TF regulators of floral development (Flower-GRN)

We hypothesize that the top TFs in the flower-GRN would be enriched with regulators of floral development. The top ten TFs ranked as the most influential in the flower-GRN were retrieved. Indeed, five well characterized floral morphology/timing regulators, namely TSO1, NAC2, NAC5, FWA and TCP22 were found in the top ten (Fig. 2B). The remaining five TFs in the top ten list (Supp. Table 4) have not yet been described as floral regulators thus providing potential candidates for testing.

#### Case study 2: Phosphate responsive genes in roots (Root-GRN)

A recent study identified the top 50 genes that are most strongly induced by phosphate supply (42). Using these genes as query we retrieved the phosphate-subnetwork from the root-GRN and identified the top 20 TFs that regulate these 50 genes (Supp. Table 5). Four known phosphate responsive regulators CPL3 (43, 44), HHO2 (45), RSL2 (46) and WRKY6 (47) are predicted to regulate these genes in the roots (Fig. 2B), with CPL3, HHO2, RSL2 and WRKY6 ranked 1^st^, 2^nd^, 4^th^ and 11^th^ respectively (Supp. Table 5).

#### Case study 3: Lipid biosynthesis in seeds (Seed-GRN)

Lipid biosynthesis in seeds is a process fundamental to the reproductive success of plants as well as essential for food and biofuel production. Due to its significance, the biosynthetic pathway leading to formation of fatty acids and storage lipids has been extensively characterized (48). All Arabidopsis genes associated with the GO term lipid biosynthetic process (GO:0008610) were retrieved (49) to form a query set of 343 genes (Supp. Table 6). The seed-GRN was queried with this set to retrieve the sub-network including these genes and their predicted regulators. This lipid-seed-GRN was further filtered to retain only those edges with an edge score >0.1 to retain only high-confidence edges (Supp. Table 7). The sum of weights for each TF was calculated within this high-confidence network and used to rank the relevance of TFs (Supp. Table 8) to lipid biosynthesis in the seeds. The top 20 ranked TFs included six known regulators of seed lipid biosynthesis (Supp Table 8) LEC1 (Rank 2^nd^) (19), MYB118 (3^rd^) and MYB115 (6^th^) (50), TT2 (4^th^) (32), FUS3 (8^th^) (20, 51), the master regulator WRI1 (9^th^)(13, 14, 16), MYB96 (14^th^) (23), and WRI3 (15^th^) (52).

#### Functional profiling of candidate seed lipid regulators validated their role in promoting seed lipid biosynthesis

In each of the three case studies above, multiple known regulators of the process were recovered in the top ranked TFs. Interspersed with the known TFs are unproven regulators for these well-established processes. To investigate the relevance of these predicted candidate regulators, we focused on a functional validation approach for the seed lipid TFs. We chose the top 20 ranked TFs to expand the list of candidate TFs. Among the top 20 TFs, we found two more known regulators MYB96 (14^th^) (23) and WRI3 (15^th^) (52), while a role for the remaining 12 candidate TFs in seed oil production has never been described.

To estimate the net effect of the perturbation of each of these TFs on lipid biosynthesis, the subset of lipid biosynthesis genes that are predicted to be regulated by these 20 TFs were placed in the context of their specific metabolic pathway using the KEGG database (Fig. 3). This approach revealed that 100 genes associated with the lipid biosynthesis process and regulated by these 20 TFs are involved in the synthesis of fatty acids (FAs) from Acetyl-CoA as well as downstream processes such as generation of Triacylglyderides (TAG), membrane lipids, steroid and terpenoid biosynthesis (Fig. 3). In the seeds, TAGs are the primary form of storage lipids and their biosynthesis occurs partially in the plastids and partially in the cytosol (11). The seed lipid GRN predicts a role for these 20 TFs in the regulation of genes involved in FA synthesis and elongation in the plastids and the formation of TAGs in the cytosol (Figure 3). For example, the gene KASIII encodes a plastid-localized enzyme with keto-acyl synthase activity that catalyzes the first step in creation of the lipid carbon chain. The seed-GRN predicts that EIL5, MybS2, TGA4, SPL12 and DiV2 regulate this crucial step in synthesis of long chain FAs. In addition to regulating TAG biosynthesis, the GRN predicts how these TFs regulate alternate metabolic fates of fatty acids and/or Acetyl-CoA (Figure 3, Supp Table 7). Of the unvalidated regulators, EIL5, bHLH93, MYB30, MybS2, CAMTA6, HB25, TGA4, DAG2, SPL12, DiV2 and AGL18 are predicted to regulate genes in one or more of these pathways. Based on these predictions, we hypothesized that lowering/eliminating the expression of any one of these eleven TFs would result in a significant reduction of total lipid content in the Arabidopsis seeds.

**Figure 3.**
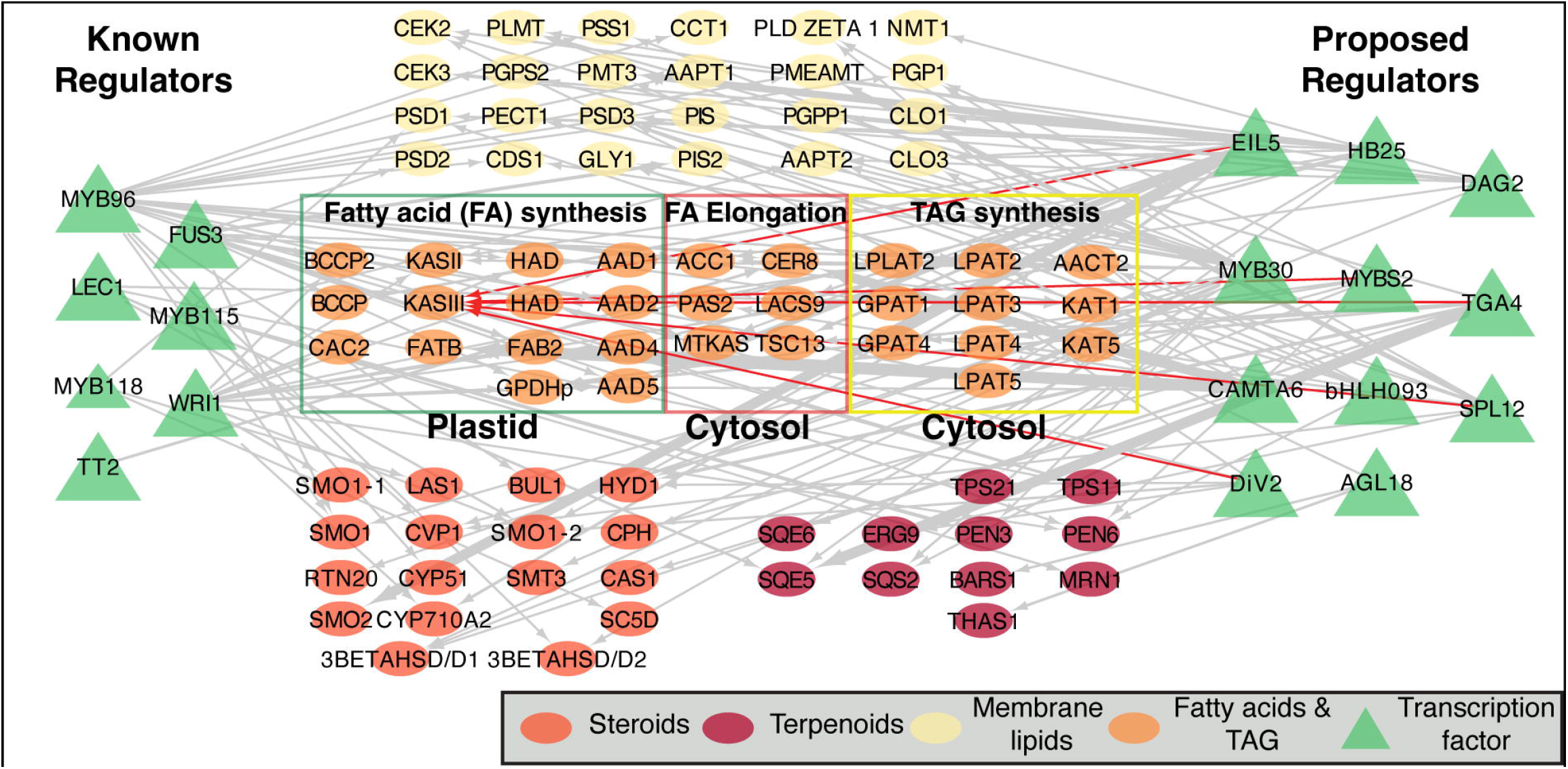
Top 20 TFs of seed lipid biosynthesis in the seed-GRN are predicted to regulate multiple steps in the lipid biosynthesis pathway. Shown here are multiple enzymes involved in the biosynthesis of membrane (yellow nodes) and storage (orange nodes) lipids as well as steroids (dark orange) and terpenoids which are fatty acid derivatives. These 106 genes are a subset of the 343 genes that were associated with the GO term lipid biosynthesis AND are assigned to one of the following pathways in the KEGG database: Glycerophoshpholipid metabolism (ath00564), fatty acid metabolism (ath01212), steroid biosynthesis (ath00100) or sesquiterpenoid and triterpenoid biosynthesis (ath00909). TFs that are predicted to regulate one or more of these genes are represented by green triangles. The TFs known to regulate the synthesis of tri-acyl-glycerides (TAGs) in seeds, are shown on the left, while the TFs predicted by the seed GRN, but not yet validated are on the right. For example, the plastid localized enzyme KASIII is predicted to be regulated by EIL5, MybS2, TGA4, SPL12 and DiV2 TFs (edges in red).

To characterize whether these candidate TFs regulate seed lipid biosynthesis as predicted by the seed-GRN, we screened the T-DNA insertion mutant lines for each of these proposed regulators to assay any changes in total fatty acid content (as fatty acid methyl esters (FAME)). Since many known seed lipid regulators also alter seed size and seed weight (14), our mutant screen additionally characterized: *i*. Seed weight (weight of 100 seeds) and *ii*. Avg. seed size (area; see Methods). We successfully isolated homozygous T-DNA insertion mutant lines for 11 of the 12 candidate TFs (Supp. Table 9) but recovered no viable mutants for EIL5. We then compared their seed phenotypes with concurrently grown WT plants (see Methods). Seed weight was significantly reduced in all tested TF mutants except *div2* and *cesta* (Fig. 4B, Supp Table 10, Supp. Figure 7). Seed size in all the candidate TF mutants except *cesta* was marginally but significantly reduced, relative to the WT seeds (Fig. 4C, Supp. Table 11, Supp. Figure 8). A significant reduction in total fatty acid (FA) content measured as FAME μg/mg of seeds, was found in five out of the 11 TFs tested: *div2*(7^th^), *mybs2*(10^th^), *tga4*(17^th^), *spl12*(19^th^) and *agl18*(20^th^) (p.adj <0.05; Fig. 4A, Supp. Table 12, Supp. Figure 6). Since some mutant lines showed a concurrent large reduction in seed weight, a constant mass (10mg) of seeds represents a very different number of seeds in these lines. Therefore, we compared the FAME per seed (see Methods) in the three mutant lines with > 20% reduction in seed weight: *myb30, hb25* and *dag2*. The per seed level of FAME is significantly reduced (p.adj <0.01) in the *myb30, hb25* and *dag2* lines (Supp. Table S12).

**Figure 4.**
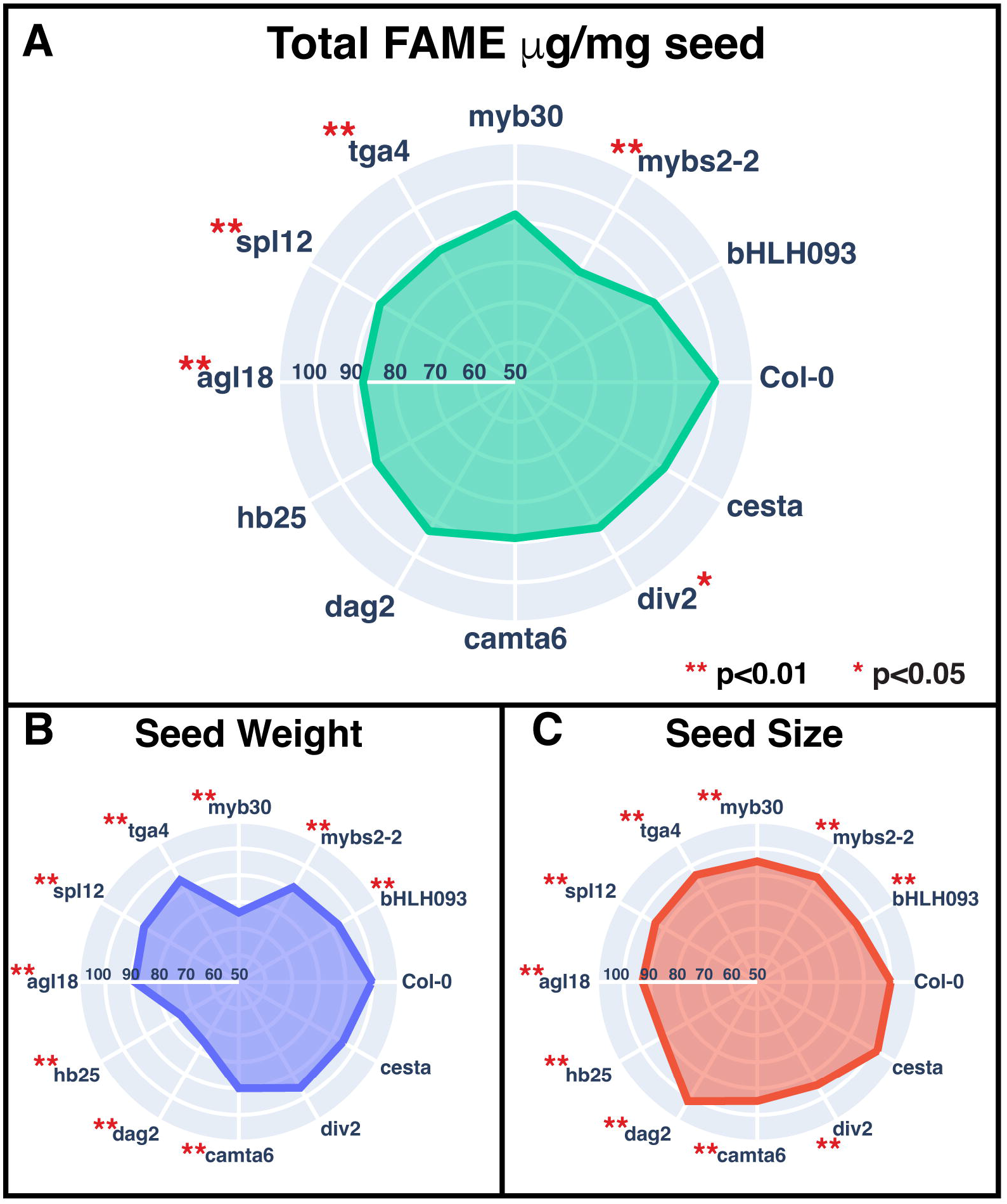
Candidate TFs show significant reduction in seed weight, size or total FA. Plotted here are the %reductions in the measured phenotype across replicates (n>=7). The WT (Col-0) is presented as 100% on the radial axis. Each inner circle represents a progressive 10% reduction from the level of the wild type (Col-0) **A**. Total FA content was significantly reduced (7-18%) in five candidate TFs with the largest reduction seen in the mybs2-2 mutant. (see also, Supp. Table 12 and Supp. Fig 6) **B**. Seed weight, measured as weight of 100 seeds, is significantly lower (6-25%) for nine candidate TFs (also Supp. Fig 7) **C**. Seed size is significantly, albeit marginally, smaller (2-9%) for nine of ten TFs shown (also Supp. Fig 8). Seed size for dag2 was 2% larger than WT.

### MybS2 promotes overall lipid production as well as the profile of fatty acid

The *mybs2-2* mutant shows the most reduction of total lipid content (∼18%; Fig. 4C, Supp. Table 12). Previous studies have established a role for MybS2 in sugar signaling (53) as well as a negative regulator of salt stress response (54). However, its role in promoting lipid biosynthesis in seeds has never been reported. To confirm the effect of MybS2 expression levels on the seed lipid biosynthesis pathway, we isolated a second T-DNA mutant (*mybs2-1*), and simultaneously generated *i*. complementation lines (*pMybS2*::*MybS2* in the *mybs2-2* background) and *ii*. overexpression lines (*p35S*::*MybS2* in WT background). Total fatty acid content in the seeds of both *mybs2-1* and *mybs2-2* mutants is significantly reduced (Fig. 5A) confirming that reduced MybS2 expression lowers seed FA content. Importantly, in the complementation lines the reduction in total fatty acid levels is rescued and is not significantly different or higher than the WT levels (Supp. Table 12, Supp. Figure 9). In the seeds of multiple independent MybS2 overexpression lines, we observed an increased accumulation of total fatty acids (Fig. 5B) and this increase is correlated with the fold change in expression level of MybS2 (r=0.7, Fig. 5C). Collectively, these results support an essential role of MybS2 in promoting seed lipid biosynthesis. To investigate whether MybS2 affects specific types of fatty acids, or just the total amount of fatty acids, we compared the proportion of the five major species of fatty acids (palmitic acid C16:0, oleic acid C18:1, linoleic acid C18:2, linolenic acid C18:3, and eicosenoic acid C20:1) among the *mybs2* mutants, MybS2 overexpression lines, and WT. Interestingly, a significant decrease in the percentage of C18:1 and an increase in C18:3 were observed in both *mybs2* mutants compared to WT (Supp. Table 12). In contrast, an increase of C18:1 and a decrease of C18:3 was detected in multiple overexpression lines (Supp. Table 12). Moreover, the weight and size of seeds were also reduced in the mutants and increased in the overexpression lines (Supp. Table 10 & 11, Supp. Figure 10). Overall, our results confirm that MybS2 positively regulates total seed lipid content, and a shift in lipid composition from C18:3 to C18:1 and seed size and seed weight.

**Figure 5.**
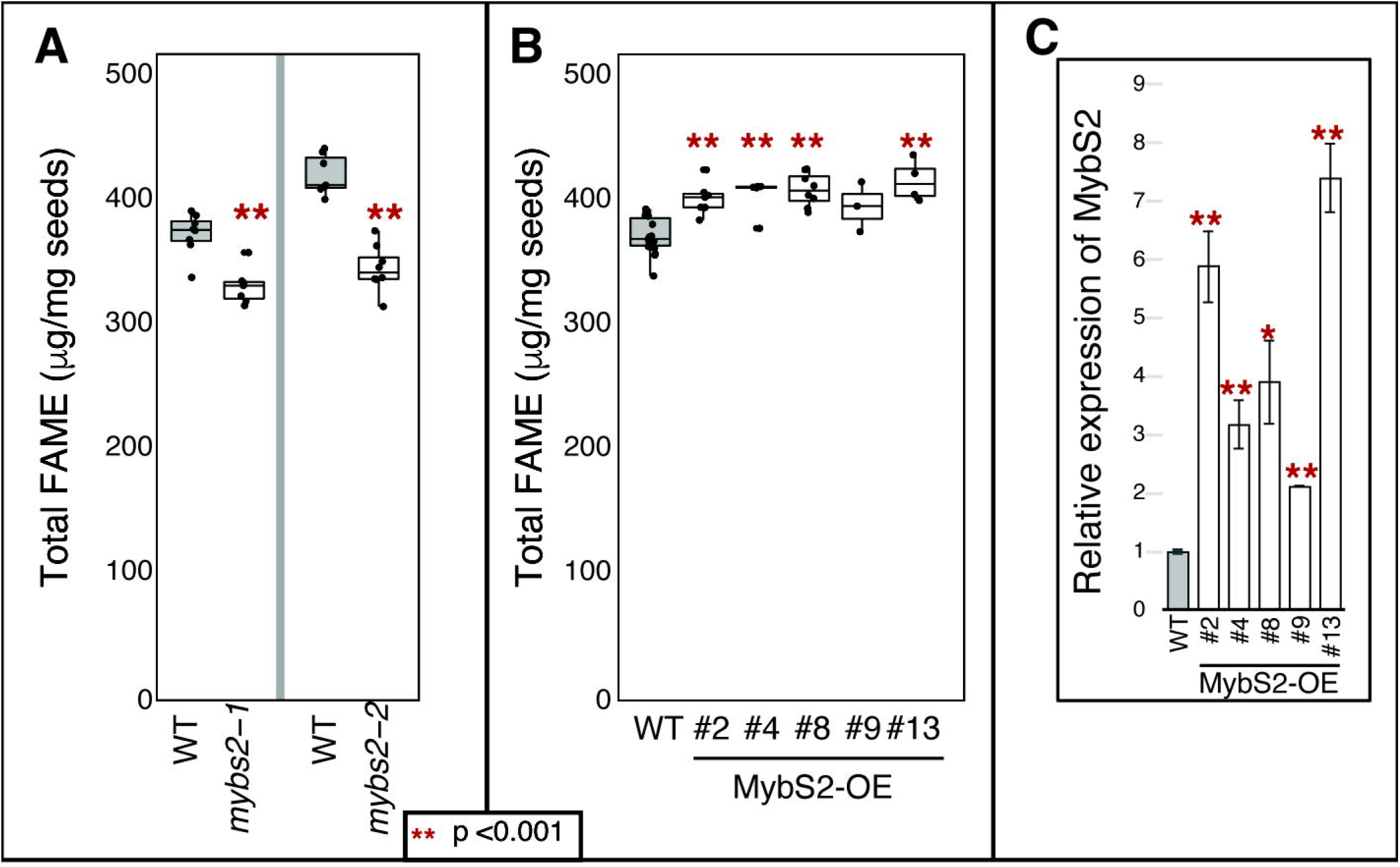
Seed oil content positively correlates with the expression level of MybS2. **A**. Total FAME was reduced by 11% in mybs2-1 and 18% in mybs2-2; two independent transposon insertion mutant lines of mybs2 **B**. Overexpression of MybS2 significantly increased the total FAME in seeds in multiple independent transgenic lines. **C**. percantage increase in Total FAME in the MybS2-0E lines correlates well (r=0.7) with the increase in expression level of MybS2 in these over-expression lines.

## DISCUSSION

This study demonstrates the benefit of building large organ delimited expression datasets. The KNN-based ML model developed here is both robust (high recall) and accurate (high precision) for every organ that had >100 gene expression datasets in the training phase. Building such organ specific gene expression sets is instrumental to GRN inference, as done here, but also to many other approaches such as correlation analysis, limiting candidate genes in QTL/GWAS analysis (e.g., expressed in the appropriate organ/correlated to known pathway genes etc.), marker gene identification, and developing organ specific promoters for synthetic biology. We applied this approach to the model plant Arabidopsis, but many other crop species such as corn, soybean, rice, tomato, wheat etc. now have public gene expressions sets with tens of thousands of gene expression assays. The ML organ classifier can be retrained for each such species to attain highest specificity. Alternately, a single classifier can be trained in one species and may predict samples across multiple plant species, as was demonstrated for prediction of cold responsive genes across species (55). The latter model would be useful even in data-poor species that lack rich expression datasets.

We built OD-GRNs for all major organs in Arabidopsis and show with multiple case studies that these GRNs are highly effective (Fig. 2B) at *de novo* identification of candidate TFs for diverse processes. Crucially, the computationally rigorous steps of OD-GRN inference are already done, so identifying candidates TFs for new processes is a relatively easy lookup, thus democratizing candidate TF discovery. Further, our study demonstrates that the candidate TFs have a high validation rate. For example, of the top twenty candidate TFs for seed lipid biosynthesis, eight were previously validated and we validate a further five TFs yielding a net success rate of 65% across the top ranked candidates. This success rate jumps up to 75% if MYB30, HB25 and DAG2 are considered validated based on the FAME/seed measure. The observed success rate is no doubt helped by the fact that the seed lipid biosynthesis process is extremely well characterized. This large and well-integrated set of genes form an ideal bait to recover TFs that regulate multiple steps of the pathway. The effect of the loss of each TF mutant on the total FA content ranged between 5-18%, which indicates that these TFs are not master regulators of seed development whose deletion is as detrimental as WRI1, FUS3, LEC1 etc. Rather, these TFs have subtler effects on lipid biosynthesis that is more similar to Myb96 (23) and Myb115/118 (50). Nonetheless, the 8-12% increase in FA content, and the 18:3 -> 18:1 compositional change observed in the MybS2-OE lines imply that genetic manipulation of these TFs, either singly or in combinations is a viable strategy to improve FA content or composition. The best strategy to engineer such increases can be designed with further knowledge of the specific lipid pathway targets for each TF as determined by further expression and TF-binding studies.

Other processes that may fit this profile include specialized metabolism pathways, rapid responses to environmental cues such as nutrients/temperature etc. and developmental processes such as organ formation. Conversely, poorly characterized processes with many unknown genes, or other genesets with poor functional coherence may indeed prove to be weaker network baits. Nonetheless, in species with existing mutant populations, or where candidate TFs can be quickly knocked out by CRISPR-based methods a reverse genetics strategy could be used to rapidly screenand validate 10-20 candidate TFs, as we have shown in this study. In the case of seed oil biosynthesis, GRN analysis permits the discovery of novel regulators from abundant expression data that may be difficult to identify through genetic screening approaches.

The accuracy of inferred GRNs are typically measured as how many of the predicted edges are known or validated (9, 10, 39, 56). The known edges are often used to calculate precision and recall (PR) thresholds and then used to cull false positives and select high-confidence edges. The relatively small subset of edges retained in this approach select for strong effects of a TF on a target gene but may potentially miss meaningful but weaker effects of a TF across multiple targets. Our approach uses a relatively lax threshold and retains the top 10% ranked edges in each OD-GRN, which no doubt includes many false predictions of TF-target interactions. Instead, we assume that the sum of weights of all edges from a TF captures the net effect of a TF on the organ or a selected process by accumulating weak effects of the TF on a broader set of targets. This approach is inspired by the concept of hidden support in phylogenomics where a tree topology unsupported by individual gene trees may derive support from concatenated matrices (57). Similarly, a TF that has weak edges to many genes involved a given process would receive a high rank even though none of its individual edges would pass a PR threshold. The three case studies presented here demonstrate the robustness of this approach.

Overall, this study describes a general approach that begins with an unfiltered gene expression dataset, which is used to build organ-delimited expression sets and infer their GRNs. The GRNs can then be used for candidate TF selection for any process of interest. This general approach is broadly applicable and across any plant and animal species that have large expression datasets.

## Materials and methods

### Assignment of organ labels for gene expression dataset

Raw sequencing data from all Illumina sequencing based gene expression data (RNASeq) from the species *Arabidopsis thaliana* were downloaded from NCBI’s SRA database. Text descriptions associated with each dataset were manually read to assign the correct organ label to each RNASeq dataset. Sequence data was converted to gene expression counts using a quasi-mapping approach (33) and normalized using a quantile normalization approach (34) and then transformed to a log2 scale.

### Machine learning pipeline to classify the organ origin of transcriptomes

A KNN-based model was trained using the scikit-learn package (58) using a minimum of 80% of the labeled samples from each organ. The maximum samples from any organ was set to 1,000 in the training set resulting in a total training set of 4,241 samples. The remaining samples of 11,010 were used as the test set. Precision and accuracy of the classifier were estimated from the test set.

### Network inference and TF ranking

For each organ, the expression set was built from the samples labeled as belonging to that organ by the multiclass classifier. A known set of transcription factors (41) in the Arabidopsis genome (regulators) and normalized expression of all genes were provided as input to the GENIE (40) inference algorithm. The top 10% ranked edges from each GRN were retained for further analysis. For any given organ or subnetwork, the edge weights were summed for each TF and this cumulative weight sum was used to rank the relative importance of TFs.

### Plant Materials and Growth Conditions

All T-DNA mutant lines were ordered from ABRC. Homozygous insertion of T-DNA was confirmed by PCR-based genotyping with LBb1.3 and Salk line-specific LP primer pairs (Supp. Table 9). Each Arabidopsis plant was grown in a 4’’ pot filled with standard germination mix (3:1) under 16/8 h light/dark conditions at 22°C in the growth room.

### Plasmid preparation and Arabidopsis transformation

To generate the *MybS2*overexpression construct the coding DNA Sequence (CDS) of the *MybS2* gene was amplified by specific primer pairs (supplementary table) using ABRC plasmid (U19230) as a template. The overexpression construct was prepared by cloning the *MybS2* CDS into the pENTR-D/TOPO vector (Invitrogen) and then mobilized to the binary vector pGWB614 (RIKEN, Japan) using Gateway™ LR Clonase™ II Enzyme mix (Thermo-Fisher Scientific). Sanger sequencing was used to confirm the orientation and frame of insert. *35S::MybS2-6HA* construct was transformed into the Arabidopsis Col-0 plant through the Agrobacterium (strain GV3101) using the floral dip method (59). Transformants (T1) were screened by spraying BASTA (Glufosinate Ammonium, 0.01%) as well as PCR with vector-specific primers (Supp. Table 14). Single insertion, T3, homozygous lines were obtained by analyzing the segregation pattern of the BASTA resistance gene. Level of expression was checked via qPCR using gene-specific primers (Supp. Table 14) with the Verso SYBR Green 1-Step qRT-PCR ROX Mix following manufacturer’s instructions. Results were calculated using the 2^−ΔCt^ method with *UBQ10* (AT4G05320) gene as an internal control. Three biological replicates of each line were used.

### Seed Phenotyping

Seed size measurement was performed as previously described (60). Seeds (1 mg) of each genotype were spread on the glass slide and photographed using a dissection microscope. The surface area of each seed was measured using ImageJ software. Seed weight was measured using an electronic balance after counting 100 seeds from each genotype manually. At least eight independent biological replicates were used for each measurement and Student’s *t*-test was used for significance evaluation.

### Total fatty acid estimation

Fatty acid extraction and analysis were performed as previously described (61). Briefly, 10 mg of Arabidopsis seeds were transmethylated in a glass vial at 90°C for 90 min in 0.3 ml of toluene and 1 ml of 5% H2SO4 (v/v methanol). Heptadecanoic acid (C:17) was used as an internal standard. After transmethylation, 1.5 ml of 0.9% NaCl solution was added and extraction was performed using 2 ml of n-hexane. Fatty Acid Methyl Esters (FAMEs) were analyzed using TriPlus RSH autosampler and Trace 1310 gas chromatography (GC) system having a 50 m x 0.25 mm FAME GC column of film thickness 0.25 um (Agilent Technologies, Santa Clara, CA) and coupled to a TSQ 8000 mass spectrometer (MS) (Thermo Fisher Scientific, Waltham, MA). A. The GC carrier gas was helium with a 1.0 ml/min linear flow rate. The programmed GC temperature gradient was as follows: time 0 minutes, 80°C then ramped to 175°C at a rate of 13°C/minute with a 5-minute hold, then ramped to 245°C at a rate of 4°C/minute with a 2 minute at the end. The GC inlet was set to 250°C and samples were injected in split mode using a split ratio of 20. The MS transfer line was set to 250°C and the MS ion source was set to 250°C and used EI ionization with 70eV. MS data were collected in scanning mode with a range of 50-500 amu. All data were analyzed with Thermo Fisher Chromeleon (Version 7.2.9) software. At least eight independent biological replicates were used for each measurement and a student t-test was used for significance evaluation. To estimate the FAME per seed, the no. of seeds per mg was estimated from the average seed weight of 100 seeds (Supp. Table S10). This estimated number of seeds per mg was then used to convert FAME μg/mg seeds to FAME/seed.

## Acknowledgement and funding sources

This material is based upon work supported by the U.S. Department of Energy, Office of Science, Office of Biological and Environmental Research (BER), under Award Number DE-SC0020399 to K.V., Y.L and K.H. This research is also supported by USDA National Institute of Food and Agriculture Hatch project numbers 1013620 to Y.L.

## Data Availability

All the data, models and networks constructed in this study will be available via links at https://www.purdue.edu/hla/sites/varalalab/infernet/. Primary gene expression data and OD-GRNs will be made available via the archival resource: Purdue University Research Repository (PURR).

## Notes

### Competing Interest Statement

The authors have declared no competing interest.

## References

1. M. Saint-Antoine, A. Singh, Network inference in systems biology: recent developments, challenges, and applications. Curr. Opin. Biotechnol. 63, 89–98 (2020).

2. S. Haque, J. S. Ahmad, N. M. Clark, C. M. Williams, R. Sozzani, Computational prediction of gene regulatory networks in plant growth and development. Curr. Opin. Plant Biol. 47, 96–105 (2019).

3. R. C. O’Malley, et al., Cistrome and epicistrome features shape the regulatory DNA landscape. Cell 166, 1598 (2016).

4. M. D. Brooks, et al., ConnecTF: A platform to integrate transcription factor–gene interactions and validate regulatory networks. Plant Physiol. 185, 49–66 (2020).

5. O. Wilkins, et al., EGRINs (Environmental Gene Regulatory Influence Networks) in Rice That Function in the Response to Water Deficit, High Temperature, and Agricultural Environments. Plant Cell 28, 2365–2384 (2016).

6. M. Schmid, et al., A gene expression map of Arabidopsis thaliana development. Nat. Genet. 37, 501–506 (2005).

7. F. F. Aceituno, N. Moseyko, S. Y. Rhee, R. A. Gutiérrez, The rules of gene expression in plants: organ identity and gene body methylation are key factors for regulation of gene expression in Arabidopsis thaliana. BMC Genomics 9, 438 (2008).

8. A. V. Klepikova, A. S. Kasianov, E. S. Gerasimov, M. D. Logacheva, A. A. Penin, A high resolution map of the Arabidopsis thaliana developmental transcriptome based on RNA-seq profiling. Plant J. 88, 1058–1070 (2016).

9. K. Varala, et al., Temporal transcriptional logic of dynamic regulatory networks underlying nitrogen signaling and use in plants. Proc. Natl. Acad. Sci. U. S. A. 115, 6494–6499 (2018).

10. M. A. Reynoso, et al., Gene regulatory networks shape developmental plasticity of root cell types under water extremes in rice. Dev. Cell 57, 1177–1192.e6 (2022).

11. C. Kuczynski, S. McCorkle, J. Keereetaweep, J. Shanklin, J. Schwender, An expanded role for the transcription factor WRINKLED1 in the biosynthesis of triacylglycerols during seed development. Front. Plant Sci. 13, 955589 (2022).

12. Y. Yang, et al., Transcriptional regulation of oil biosynthesis in seed plants: Current understanding, applications, and perspectives. Plant Commun 3, 100328 (2022).

13. N. Focks, C. Benning, wrinkled1: A novel, low-seed-oil mutant of Arabidopsis with a deficiency in the seed-specific regulation of carbohydrate metabolism. Plant Physiol. 118, 91–101 (1998).

14. A. Cernac, C. Benning, WRINKLED1 encodes an AP2/EREB domain protein involved in the control of storage compound biosynthesis in Arabidopsis. Plant J. 40, 575–585 (2004).

15. Q. Kong, W. Ma, WRINKLED1 transcription factor: How much do we know about its regulatory mechanism? Plant Sci. 272, 153–156 (2018).

16. S. Baud, et al., WRINKLED1 specifies the regulatory action of LEAFY COTYLEDON2 towards fatty acid metabolism during seed maturation in Arabidopsis. Plant J. 50, 825–838 (2007).

17. R. Tian, et al., Direct and indirect targets of the arabidopsis seed transcription factor ABSCISIC ACID INSENSITIVE3. Plant J. 103, 1679–1694 (2020).

18. Z. Yang, et al., ABA-INSENSITIVE 3 with or without FUSCA3 highly up-regulates lipid droplet proteins and activates oil accumulation. J. Exp. Bot. 73, 2077–2092 (2022).

19. J. Mu, et al., LEAFY COTYLEDON1 is a key regulator of fatty acid biosynthesis in Arabidopsis. Plant Physiol. 148, 1042–1054 (2008).

20. M. Zhang, X. Cao, Q. Jia, J. Ohlrogge, FUSCA3 activates triacylglycerol accumulation in Arabidopsis seedlings and tobacco BY2 cells. Plant J. 88, 95–107 (2016).

21. A. Yamamoto, et al., Diverse roles and mechanisms of gene regulation by the Arabidopsis seed maturation master regulator FUS3 revealed by microarray analysis. Plant Cell Physiol. 51, 2031–2046 (2010).

22. D. Li, et al., MYB89 Transcription Factor Represses Seed Oil Accumulation. Plant Physiol. 173, 1211–1225 (2017).

23. H. G. Lee, H. Kim, M. C. Suh, H. U. Kim, P. J. Seo, The MYB96 Transcription Factor Regulates Triacylglycerol Accumulation by Activating DGAT1 and PDAT1 Expression in Arabidopsis Seeds. Plant Cell Physiol. 59, 1432–1442 (2018).

24. A. Fatihi, A. M. Zbierzak, P. Dörmann, Alterations in seed development gene expression affect size and oil content of Arabidopsis seeds. Plant Physiol. 163, 973–985 (2013).

25. K. Geilen, M. Heilmann, S. Hillmer, M. Böhmer, WRKY43 regulates polyunsaturated fatty acid content and seed germination under unfavourable growth conditions. Sci. Rep. 7, 14235 (2017).

26. A. Mendes, et al., bZIP67 regulates the omega-3 fatty acid content of Arabidopsis seed oil by activating fatty acid desaturase3. Plant Cell 25, 3104–3116 (2013).

27. G. Barthole, et al., MYB118 represses endosperm maturation in seeds of Arabidopsis. Plant Cell 26, 3519–3537 (2014).

28. G. Song, et al., The WRKY6 transcription factor affects seed oil accumulation and alters fatty acid compositions in Arabidopsis thaliana. Physiol. Plant. 169, 612–624 (2020).

29. S. Duan, et al., MYB76 Inhibits Seed Fatty Acid Accumulation in Arabidopsis. Front. Plant Sci. 8, 226 (2017).

30. L. Shi, V. Katavic, Y. Yu, L. Kunst, G. Haughn, Arabidopsis glabra2 mutant seeds deficient in mucilage biosynthesis produce more oil. Plant J. 69, 37–46 (2012).

31. M. Chen, et al., TRANSPARENT TESTA GLABRA1 Regulates the Accumulation of Seed Storage Reserves in Arabidopsis. Plant Physiol. 169, 391–402 (2015).

32. M. Chen, et al., The effect of TRANSPARENT TESTA2 on seed fatty acid biosynthesis and tolerance to environmental stresses during young seedling establishment in Arabidopsis. Plant Physiol. 160, 1023–1036 (2012).

33. R. Patro, G. Duggal, M. I. Love, R. A. Irizarry, C. Kingsford, Salmon provides fast and bias-aware quantification of transcript expression. Nat. Methods 14, 417–419 (2017).

34. D. Risso, K. Schwartz, G. Sherlock, S. Dudoit, GC-content normalization for RNA-Seq data. BMC Bioinformatics 12, 480 (2011).

35. C.-Y. Cheng, et al., Araport11: a complete reannotation of the Arabidopsis thaliana reference genome. Plant J. 89, 789–804 (2017).

36. N. Sapoval, et al., Current progress and open challenges for applying deep learning across the biosciences. Nat. Commun. 13, 1728 (2022).

37. J. G. Greener, S. M. Kandathil, L. Moffat, D. T. Jones, A guide to machine learning for biologists. Nat. Rev. Mol. Cell Biol. 23, 40–55 (2022).

38. I. Yanai, et al., Genome-wide midrange transcription profiles reveal expression level relationships in human tissue specification. Bioinformatics 21, 650–659 (2005).

39. D. Marbach, et al., Wisdom of crowds for robust gene network inference. Nat. Methods 9, 796–804 (2012).

40. V. A. Huynh-Thu, A. Irrthum, L. Wehenkel, P. Geurts, Inferring regulatory networks from expression data using tree-based methods. PLoS One 5 (2010).

41. J. Jin, et al., PlantTFDB 4.0: toward a central hub for transcription factors and regulatory interactions in plants. Nucleic Acids Res. 45, D1040–D1045 (2017).

42. W.-R. Scheible, et al., Elucidating the unknown transcriptional responses and PHR1-mediated biotic and abiotic stress tolerance during phosphorus limitation. J. Exp. Bot. 74, 2083–2111 (2023).

43. R. Tominaga-Wada, T. Wada, Extended C termini of CPC-LIKE MYB proteins confer functional diversity in Arabidopsis epidermal cell differentiation. Development 144, 2375–2380 (2017).

44. M. Ohmagari, Y. Kono, R. Tominaga, Effect of phosphate starvation on CAPRICE homolog gene expression in the root of Arabidopsis. Plant Biotechnol. 37, 349–352 (2020).

45. X. Wang, et al., The Transcription Factor NIGT1.2 Modulates Both Phosphate Uptake and Nitrate Influx during Phosphate Starvation in Arabidopsis and Maize. Plant Cell 32, 3519–3534 (2020).

46. S. Mangano, S. P. Denita-Juarez, E. Marzol, C. Borassi, J. M. Estevez, High Auxin and High Phosphate Impact on RSL2 Expression and ROS-Homeostasis Linked to Root Hair Growth in Arabidopsis thaliana. Front. Plant Sci. 9, 1164 (2018).

47. Y.-F. Chen, et al., The WRKY6 transcription factor modulates PHOSPHATE1 expression in response to low Pi stress in Arabidopsis. Plant Cell 21, 3554–3566 (2009).

48. Y. Li-Beisson, et al., Acyl-lipid metabolism. Arabidopsis Book 11, e0161 (2013).

49. S. Carbon, et al., AmiGO: online access to ontology and annotation data. Bioinformatics 25, 288–289 (2009).

50. M. A. Troncoso-Ponce, et al., Transcriptional activation of two delta-9 palmitoyl-ACP desaturase genes by MYB115 and MYB118 is critical for biosynthesis of omega-7 monounsaturated fatty acids in the endosperm of Arabidopsis seeds. Plant Cell 28, 2666–2682 (2016).

51. H. Wang, J. Guo, K. N. Lambert, Y. Lin, Developmental control of Arabidopsis seed oil biosynthesis. Planta 226, 773–783 (2007).

52. A. To, et al., WRINKLED transcription factors orchestrate tissue-specific regulation of fatty acid biosynthesis in Arabidopsis. Plant Cell 24, 5007–5023 (2012).

53. Y.-S. Chen, et al., Sugar starvation-regulated MYBS2 and 14-3-3 protein interactions enhance plant growth, stress tolerance, and grain weight in rice. Proc. Natl. Acad. Sci. U. S. A. 116, 21925–21935 (2019).

54. T. Wang, et al., Salt-Related MYB1 Coordinates Abscisic Acid Biosynthesis and Signaling during Salt Stress in Arabidopsis. Plant Physiol. 169, 1027–1041 (2015).

55. X. Meng, et al., Predicting transcriptional responses to cold stress across plant species. Proc. Natl. Acad. Sci. U. S. A. 118 (2021).

56. M. D. Brooks, et al., Network Walking charts transcriptional dynamics of nitrogen signaling by integrating validated and predicted genome-wide interactions. Nat. Commun. 10, 1569 (2019).

57. R. H. Baker, R. DeSalle, Multiple sources of character information and the phylogeny of Hawaiian drosophilids. Syst. Biol. 46, 654–673 (1997).

58. F. Pedregosa, et al., Scikit-learn: Machine learning in Python. the Journal of machine Learning research 12, 2825–2830 (2011).

59. S. J. Clough, A. F. Bent, Floral dip: a simplified method for Agrobacterium-mediated transformation of Arabidopsis thaliana. Plant J. 16, 735–743 (1998).

60. R. P. Herridge, R. C. Day, S. Baldwin, R. C. Macknight, Rapid analysis of seed size in Arabidopsis for mutant and QTL discovery. Plant Methods 7, 3 (2011).

61. Y. Li, F. Beisson, M. Pollard, J. Ohlrogge, Oil content of Arabidopsis seeds: the influence of seed anatomy, light and plant-to-plant variation. Phytochemistry 67, 904–915 (2006).

