## Supplementary Figures for "Genome-wide, Organ-delimited gene regulatory networks (OD-GRNs) provide high accuracy in candidate TF selection across diverse processes"

### Slide 1
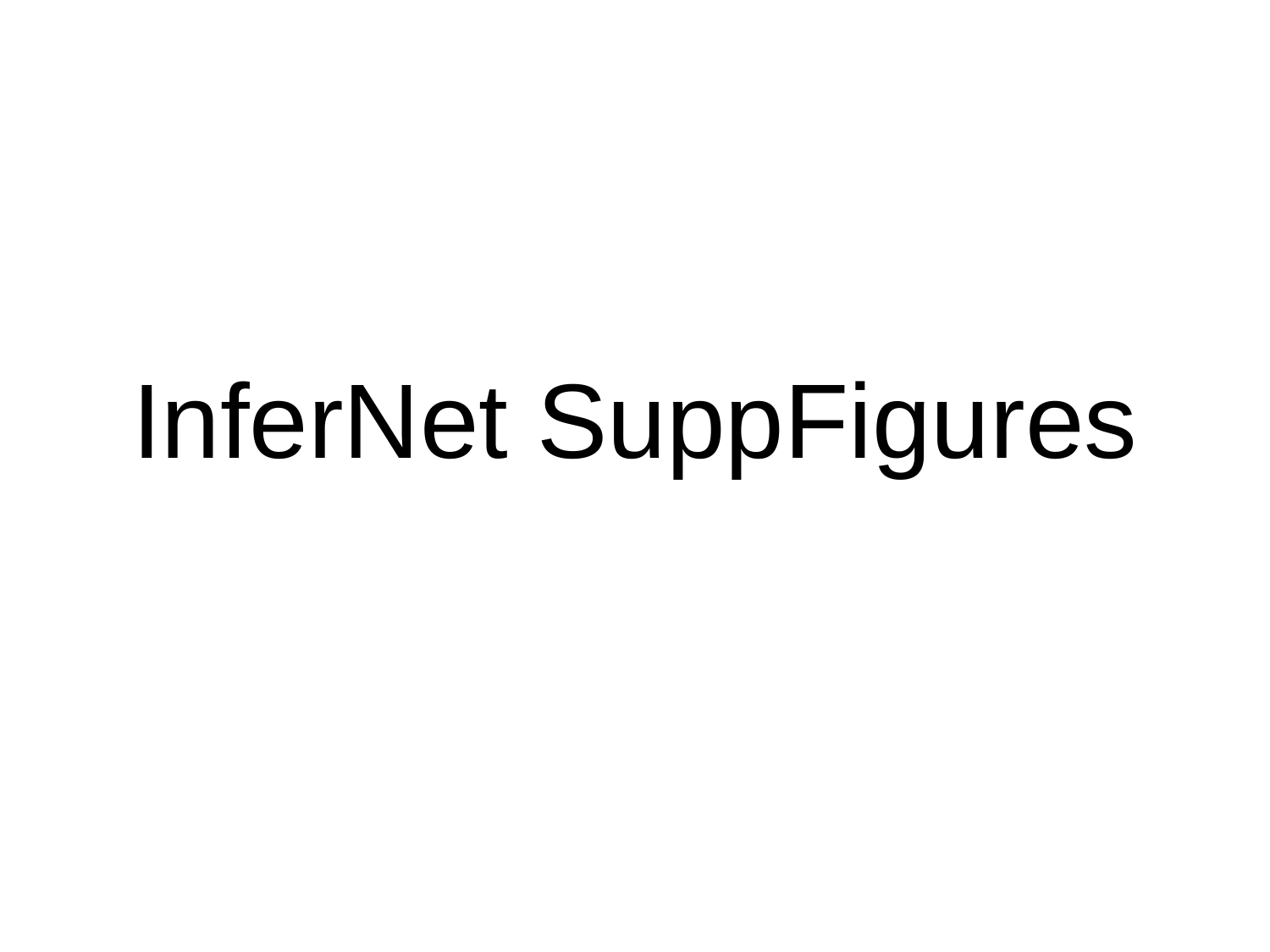

#### InferNet SuppFigures

### Slide 2
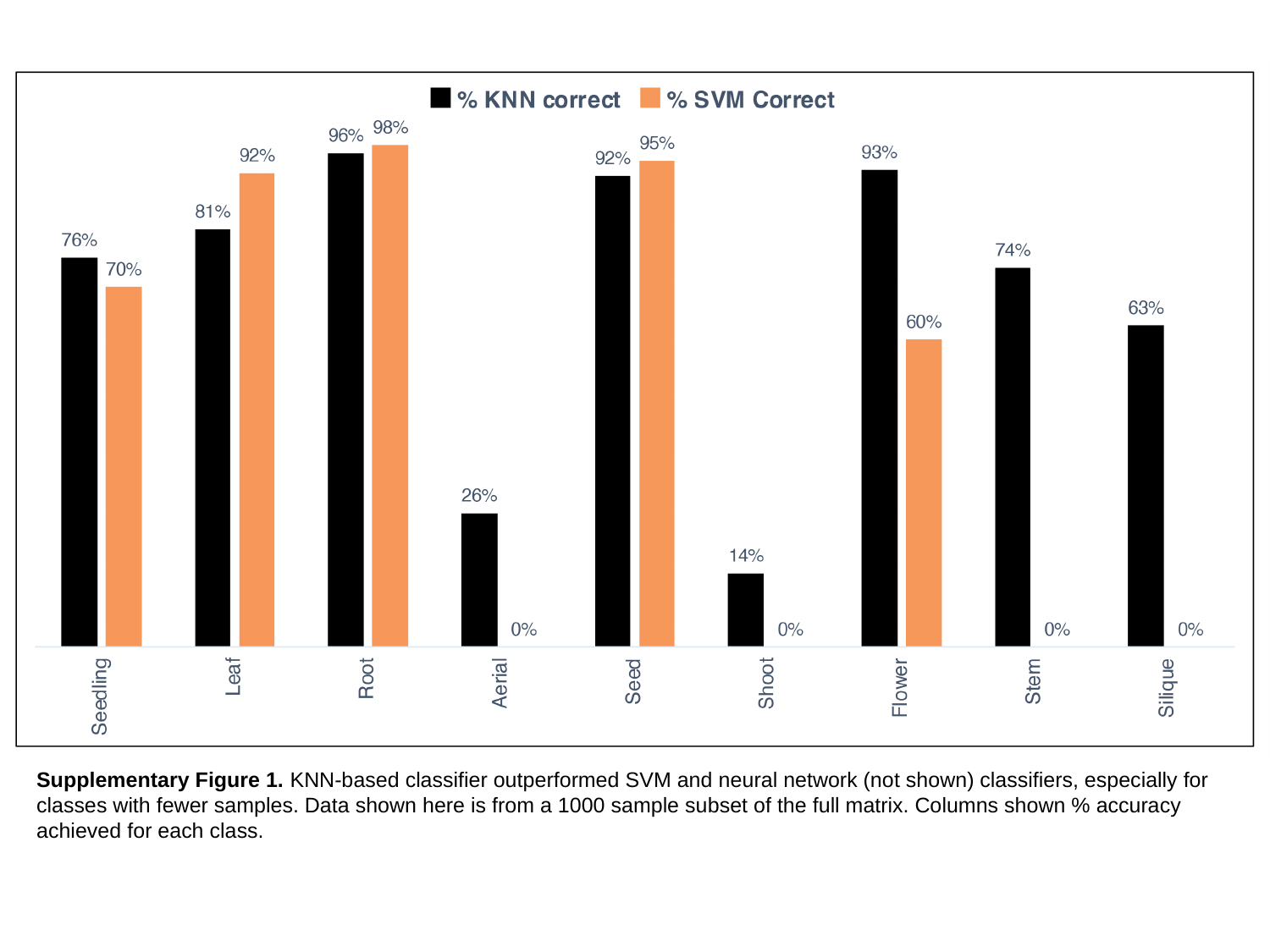

Supplementary Figure 1. KNN-based classifier outperformed SVM and neural network (not shown) classifiers, especially for classes with fewer samples. Data shown here is from a 1000 sample subset of the full matrix. Columns shown % accuracy achieved for each class.

### Slide 3
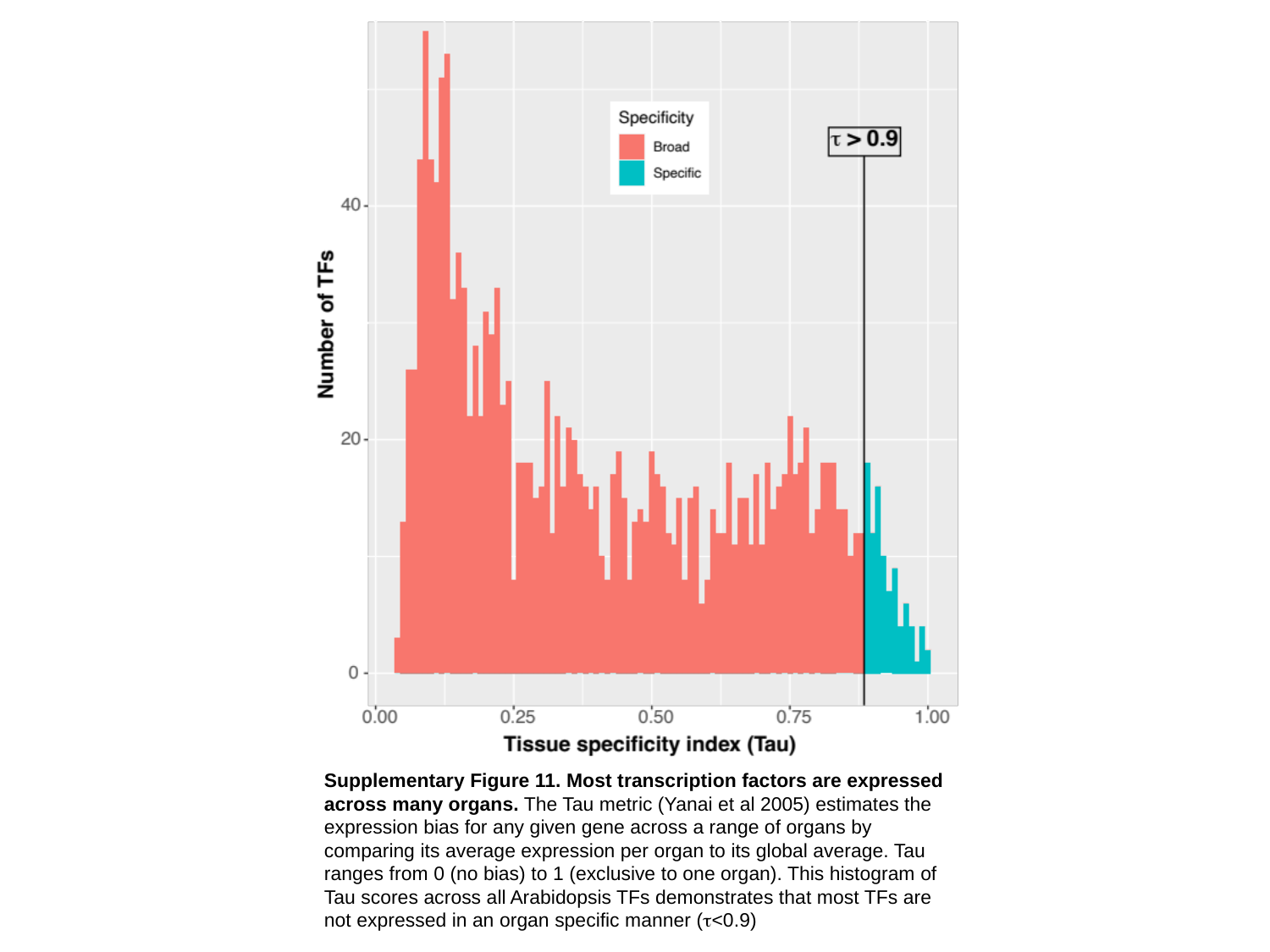

Supplementary Figure 11. Most transcription factors are expressed across many organs. The Tau metric (Yanai et al 2005) estimates the expression bias for any given gene across a range of organs by comparing its average expression per organ to its global average. Tau ranges from 0 (no bias) to 1 (exclusive to one organ). This histogram of Tau scores across all Arabidopsis TFs demonstrates that most TFs are not expressed in an organ specific manner (τ<0.9)

### Slide 4
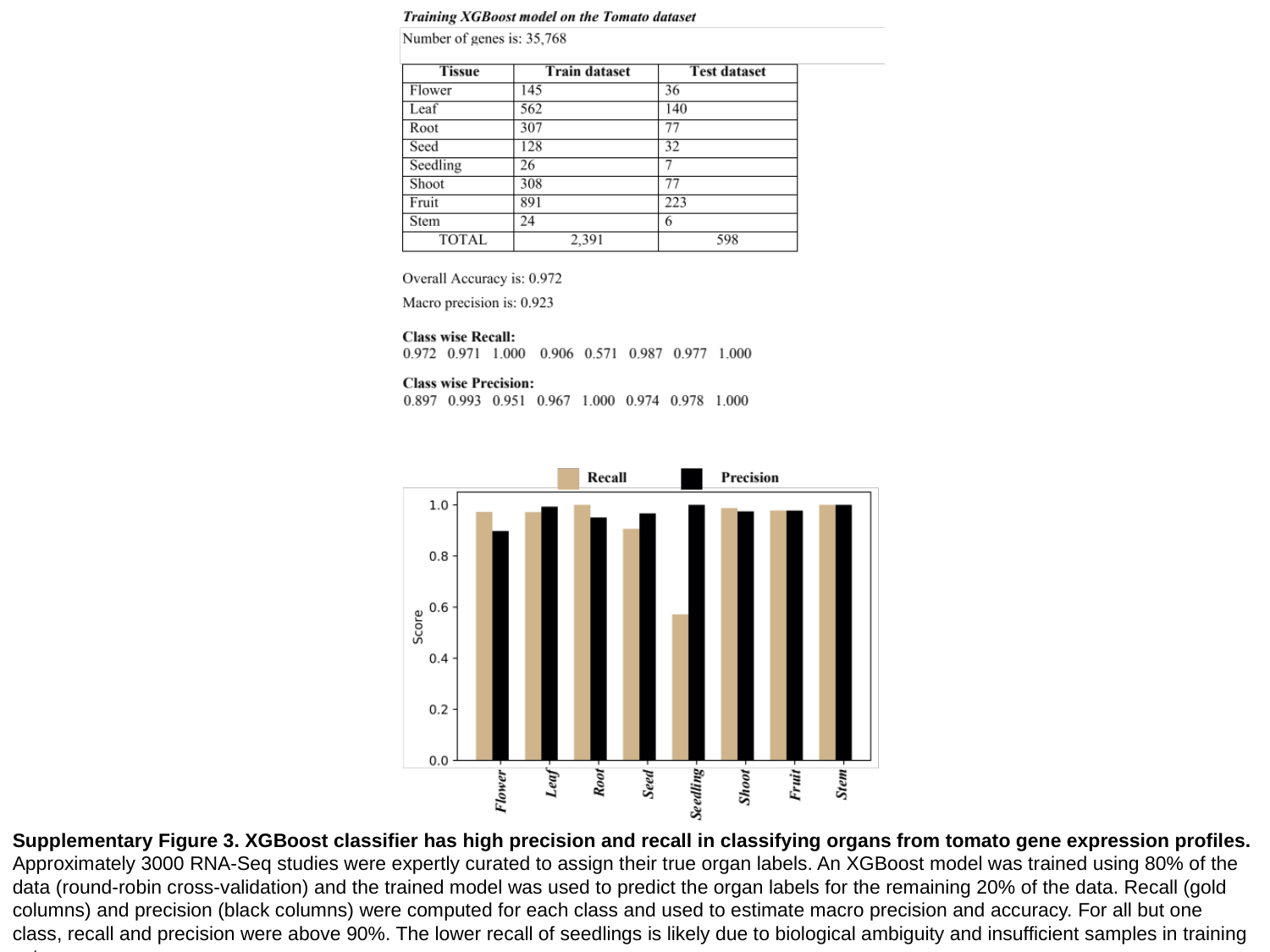

Supplementary Figure 3. XGBoost classifier has high precision and recall in classifying organs from tomato gene expression profiles. Approximately 3000 RNA-Seq studies were expertly curated to assign their true organ labels. An XGBoost model was trained using 80% of the data (round-robin cross-validation) and the trained model was used to predict the organ labels for the remaining 20% of the data. Recall (gold columns) and precision (black columns) were computed for each class and used to estimate macro precision and accuracy. For all but one class, recall and precision were above 90%. The lower recall of seedlings is likely due to biological ambiguity and insufficient samples in training set.

### Slide 5
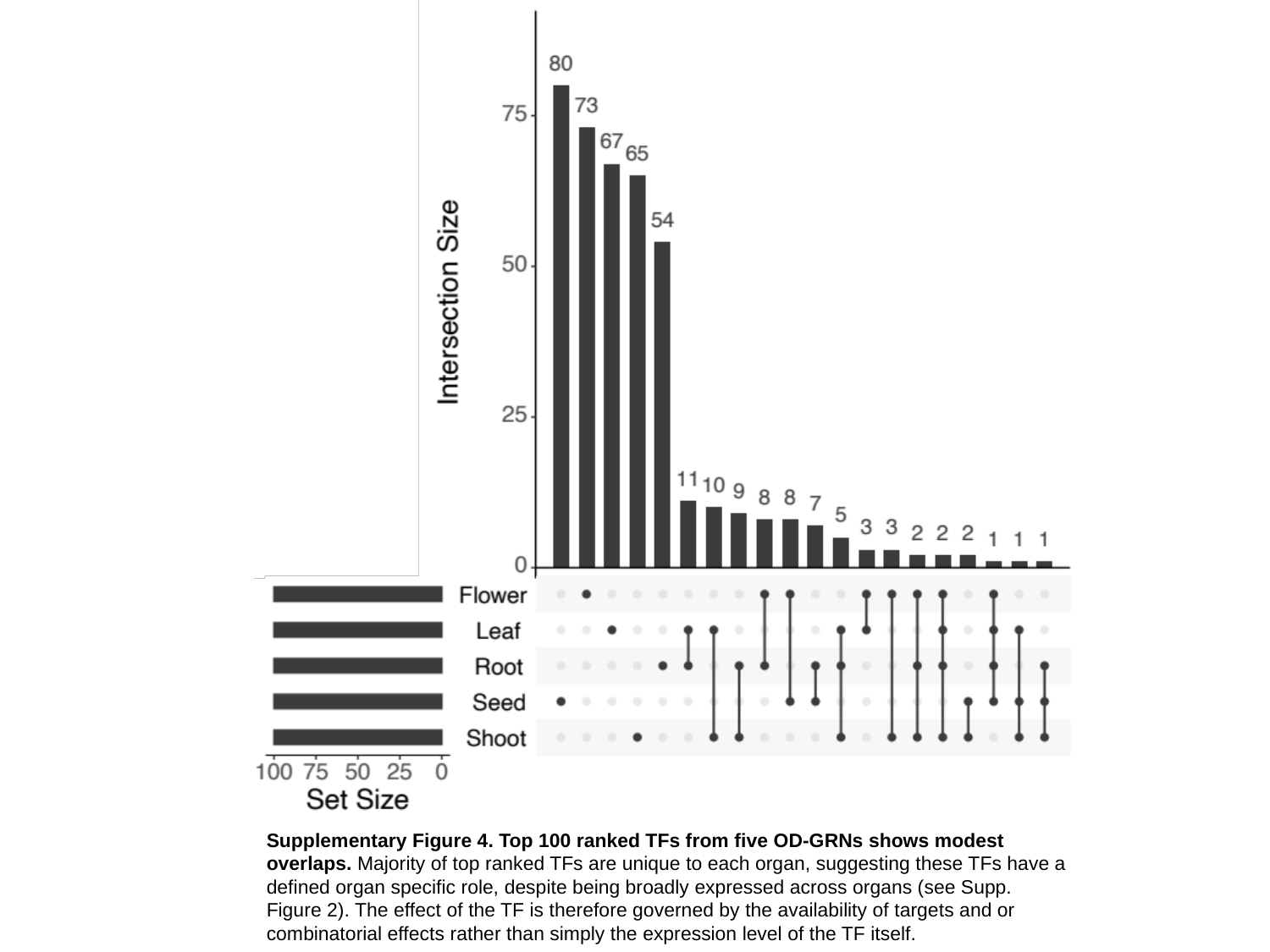

Supplementary Figure 4. Top 100 ranked TFs from five OD-GRNs shows modest overlaps. Majority of top ranked TFs are unique to each organ, suggesting these TFs have a defined organ specific role, despite being broadly expressed across organs (see Supp. Figure 2). The effect of the TF is therefore governed by the availability of targets and or combinatorial effects rather than simply the expression level of the TF itself.

### Slide 6
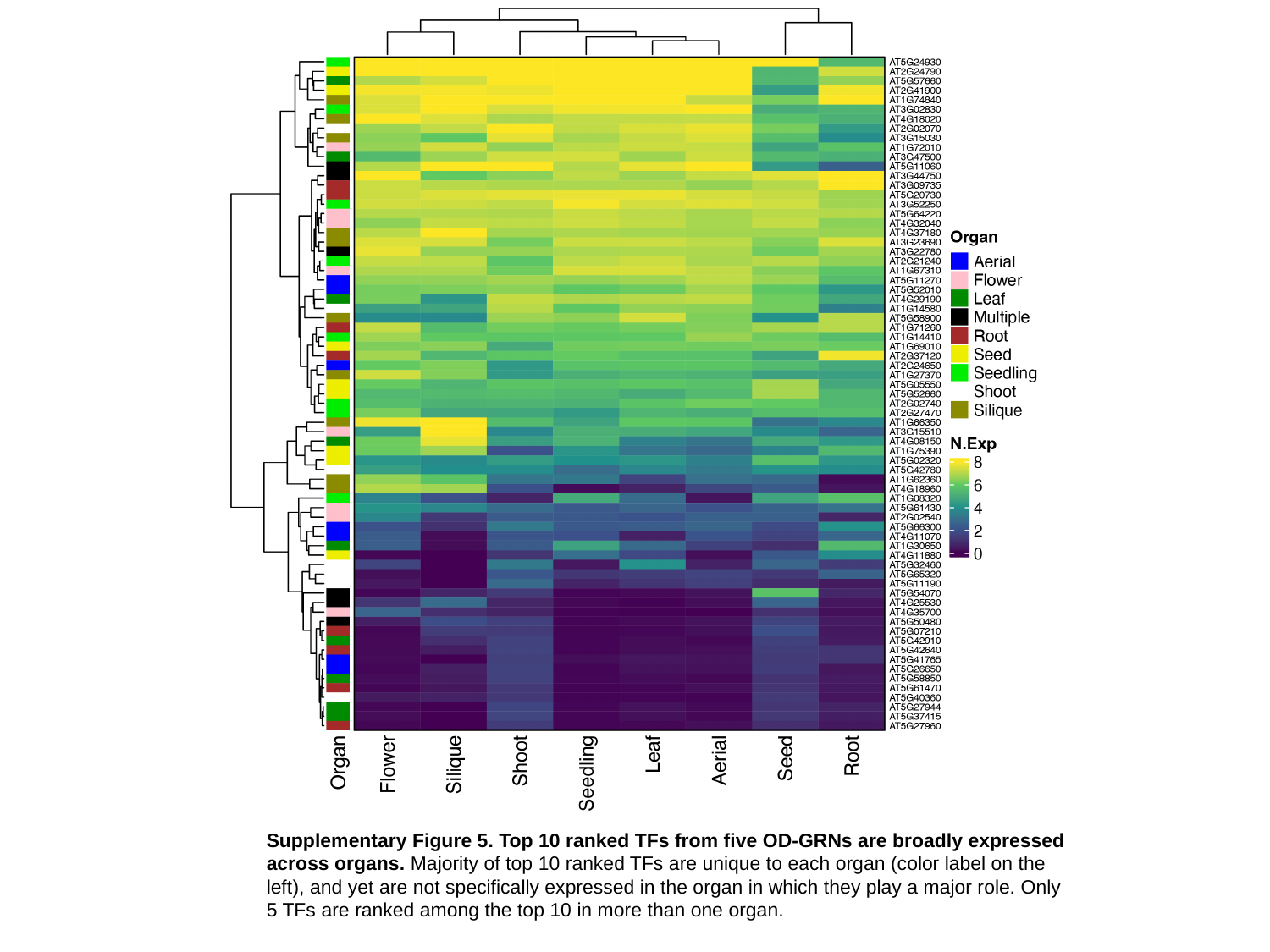

Supplementary Figure 5. Top 10 ranked TFs from five OD-GRNs are broadly expressed across organs. Majority of top 10 ranked TFs are unique to each organ (color label on the left), and yet are not specifically expressed in the organ in which they play a major role. Only 5 TFs are ranked among the top 10 in more than one organ.

### Slide 7
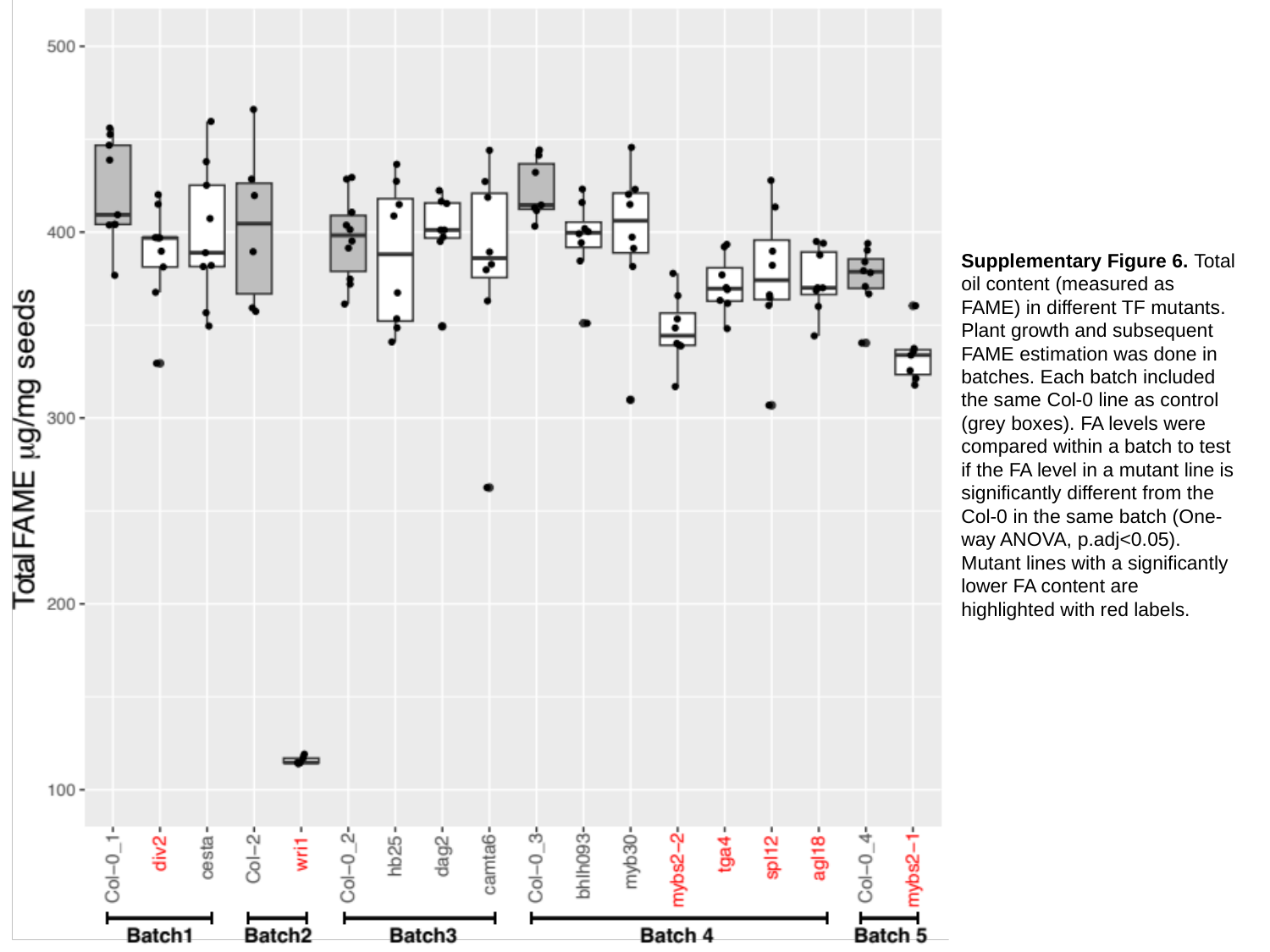

Supplementary Figure 6. Total oil content (measured as FAME) in different TF mutants. Plant growth and subsequent FAME estimation was done in batches. Each batch included the same Col-0 line as control (grey boxes). FA levels were compared within a batch to test if the FA level in a mutant line is significantly different from the Col-0 in the same batch (One-way ANOVA, p.adj<0.05). Mutant lines with a significantly lower FA content are highlighted with red labels.

### Slide 8
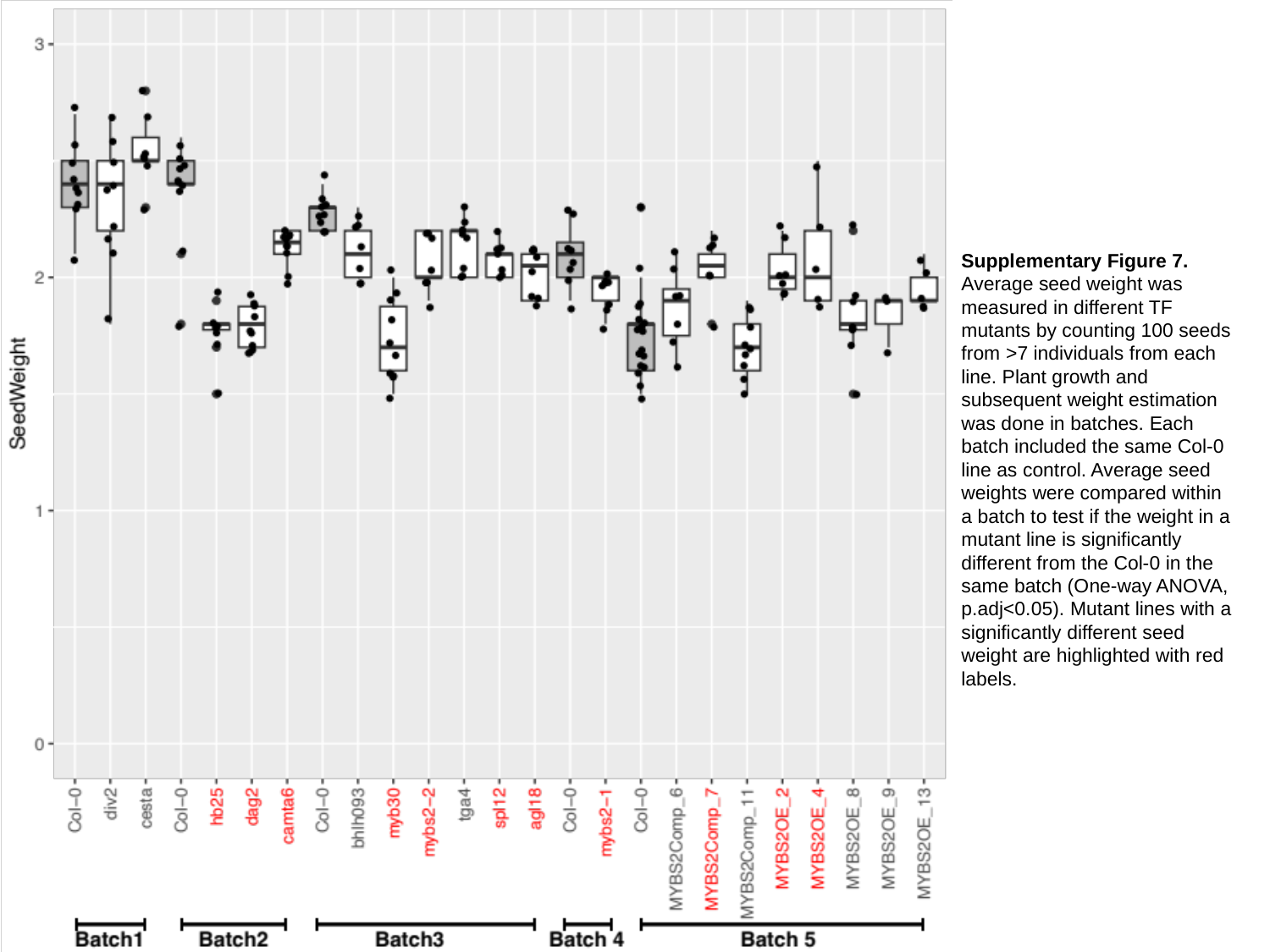

Supplementary Figure 7. Average seed weight was measured in different TF mutants by counting 100 seeds from >7 individuals from each line. Plant growth and subsequent weight estimation was done in batches. Each batch included the same Col-0 line as control. Average seed weights were compared within a batch to test if the weight in a mutant line is significantly different from the Col-0 in the same batch (One-way ANOVA, p.adj<0.05). Mutant lines with a significantly different seed weight are highlighted with red labels.

### Slide 9
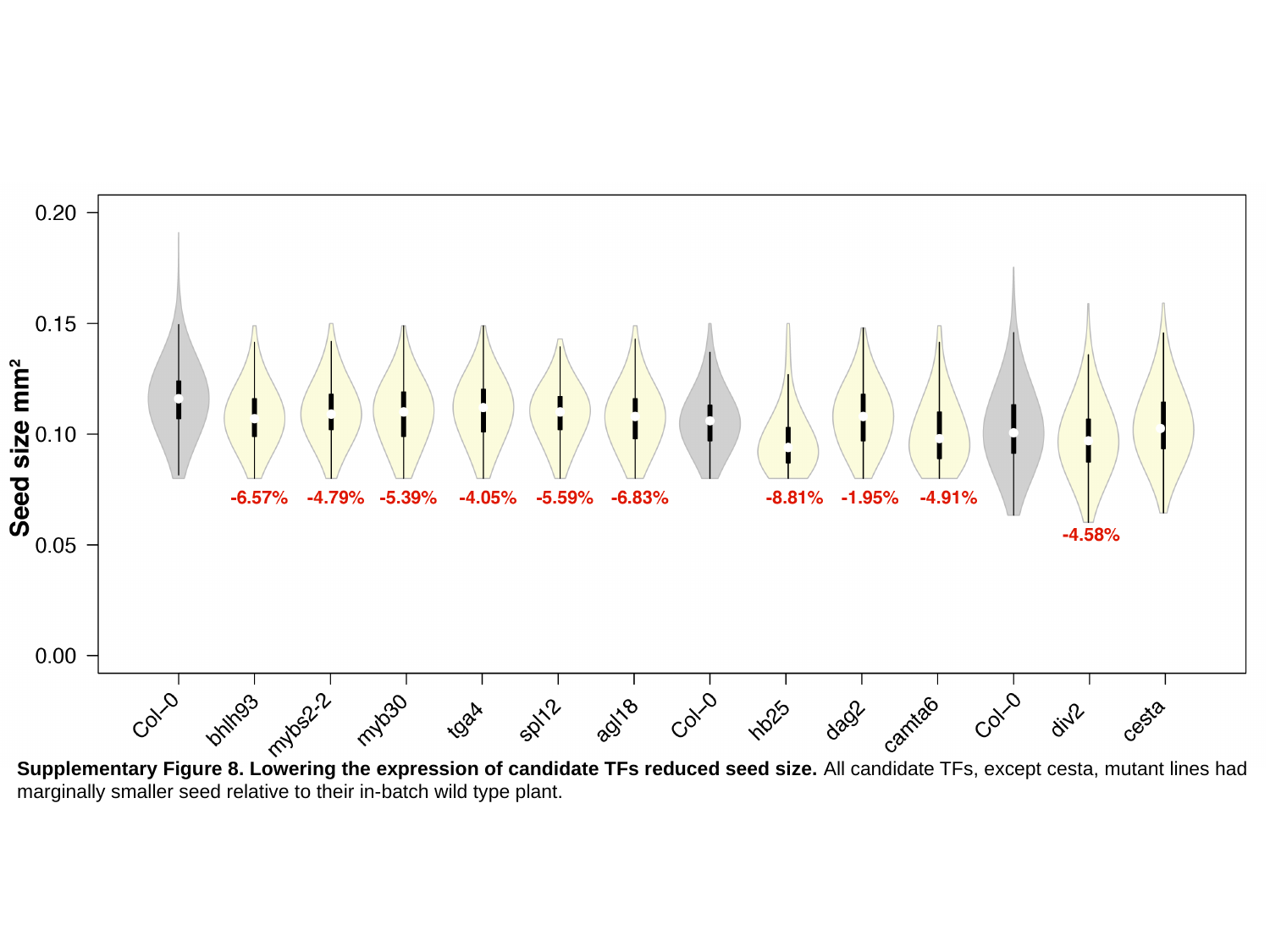

Supplementary Figure 8. Lowering the expression of candidate TFs reduced seed size. All candidate TFs, except cesta, mutant lines had marginally smaller seed relative to their in-batch wild type plant.

### Slide 10
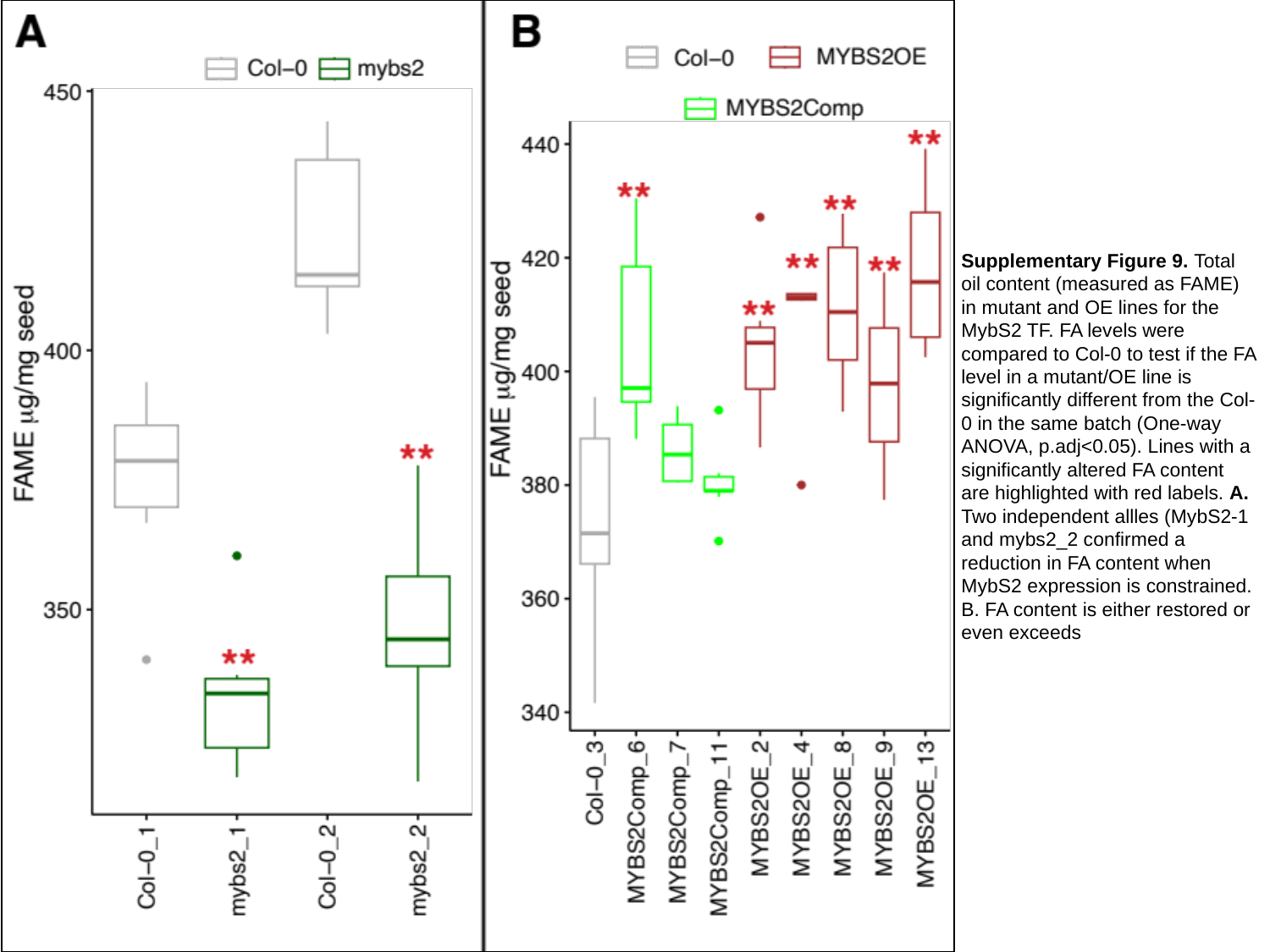

Supplementary Figure 9. Total oil content (measured as FAME) in mutant and OE lines for the MybS2 TF. FA levels were compared to Col-0 to test if the FA level in a mutant/OE line is significantly different from the Col-0 in the same batch (One-way ANOVA, p.adj<0.05). Lines with a significantly altered FA content are highlighted with red labels. A. Two independent allles (MybS2-1 and mybs2_2 confirmed a reduction in FA content when MybS2 expression is constrained.
B. FA content is either restored or even exceeds

### Slide 11
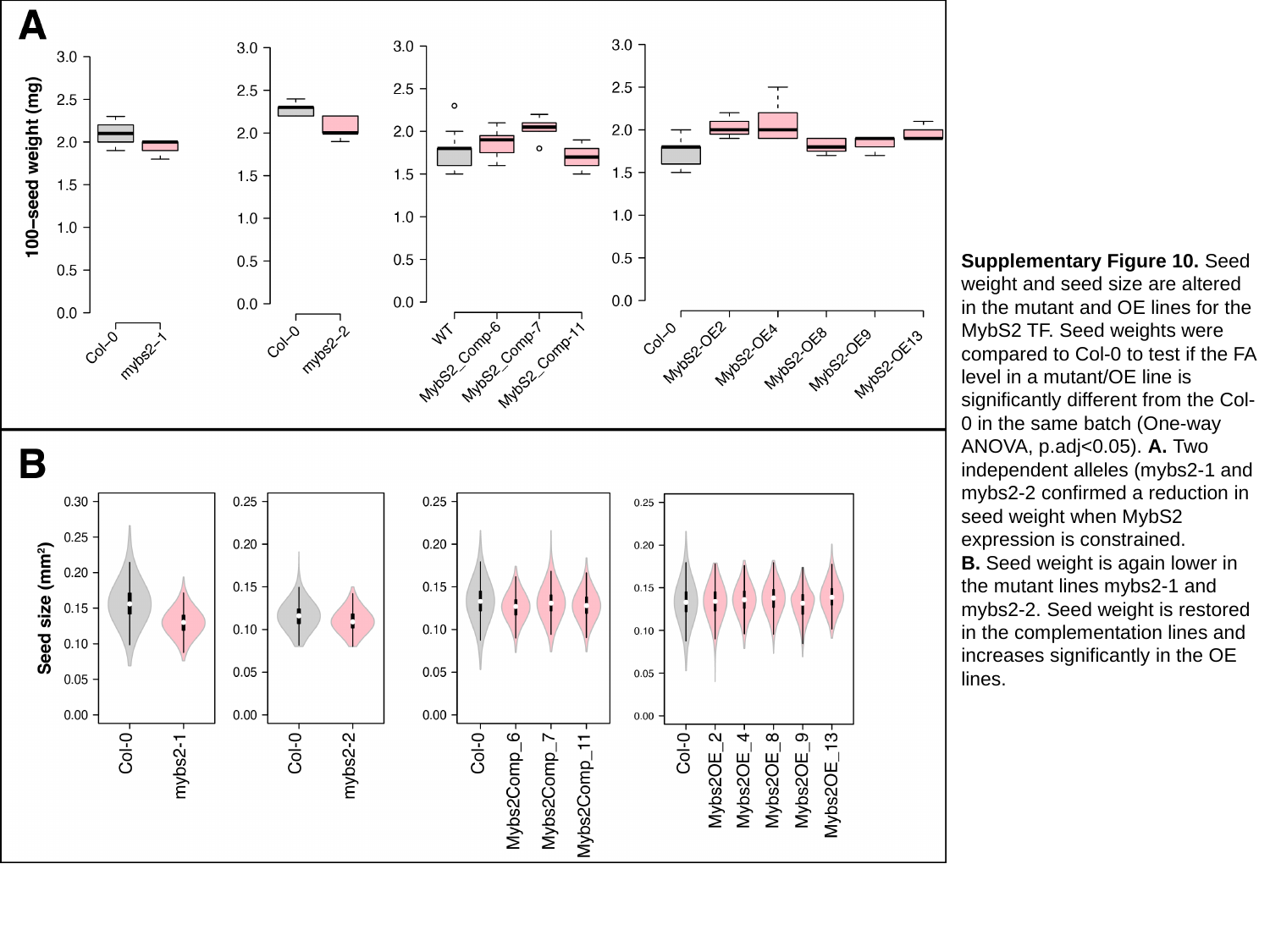

Supplementary Figure 10. Seed weight and seed size are altered in the mutant and OE lines for the MybS2 TF. Seed weights were compared to Col-0 to test if the FA level in a mutant/OE line is significantly different from the Col-0 in the same batch (One-way ANOVA, p.adj<0.05). A. Two independent alleles (mybs2-1 and mybs2-2 confirmed a reduction in seed weight when MybS2 expression is constrained.
B. Seed weight is again lower in the mutant lines mybs2-1 and mybs2-2. Seed weight is restored in the complementation lines and increases significantly in the OE lines.
